# Unifying the genetic landscape of common and rare diseases via latent neighborhoods

**DOI:** 10.64898/2026.01.27.701945

**Authors:** Diederik S. Laman Trip, Ellen Aarts, Gudmundur Magnusson, Dennis Gankin, Inigo Barrio-Hernandez, Pedro Beltrao

## Abstract

Resolving disease mechanisms and identifying safer drug targets remains challenging due to difficulties in integrating inconsistent sources of genetic evidence. Here, we developed a deep-learning (DL) method for obtaining latent representations of human diseases and other terms by coupling a variational autoencoder (VAE) to network embeddings from graph representation learning on a protein interaction network. We apply this approach to map the landscape of human disease, by systematically integrating evidence for ∼42,000 traits and terms from sources such as clinical reports (ClinVar), genome-wide association studies (GWAS), mouse phenotypes and gene ontology. Similar diseases share their latent neighborhoods with the sources of evidence highlighting different aspects of the same cell biology for its role in disease. By decoding latent neighborhoods, we unify the sources of evidence and integrate across diseases to prioritize genes for common and rare diseases. Through examples for diabetes, hearing loss, and familial long QT syndrome, we illustrate how the methodology can be applied to develop hypotheses for disease mechanisms and to propose experimental models, measurements, targets and potential drugs that may improve diagnosis and treatments.

## INTRODUCTION

Various sources of evidence link human traits and (rare) diseases to genetic variation and causal genes, such as clinical reports (ClinVar) and genome-wide association studies (GWAS). ClinVar reports variants that are primarily medically relevant and supported by evidence such as observations from clinical genetic testing (Landrum and Kattman 2018), identifying variants that are typically protein coding and with large effect sizes (Sazonovs and Barrett 2018). In contrast, GWAS link human genetic variants to traits by assaying single-nucleotide polymorphisms (SNPs) at the genome-wide scale through population-level studies (Welter et al. 2014), identifying variants that are typically non-coding, have small effect sizes, and may be mapped to likely causal genes through various approaches (Zhu et al. 2016; Mountjoy et al. 2021). Despite these differences, the diverging evidence at the variant- and gene-levels likely shows different views of shared disease mechanism(s), and these mechanisms are in turn likely similar for related diseases.

One approach to explore such shared disease mechanisms and to integrate discordant sources of evidence is through molecular interaction networks. Indeed, network-based methods are a powerful approach for uncovering disease mechanisms and prioritizing disease genes, and have been extensively applied in particular for GWAS data (Fang et al. 2019; Lee et al. 2011; Greene et al. 2015; Huang et al. 2018) and using network propagation (Barrio-Hernandez et al. 2023; Aarts et al. 2025; Leger et al. 2024) or deep learning approaches (Peng et al. 2019; Cinaglia and Cannataro 2023; Li et al. 2021; Ma et al. 2023; Hu et al. 2025). For example, such studies have explored the pleiotropy of common GWAS traits (Barrio-Hernandez et al. 2023), disease modules for related rare disorders (Buphamalai et al. 2021), the etiology of rare monogenic disorders such as ciliopathies (Aarts et al. 2025), and the shared biology for common and rare variants in the context of alcohol consumption (Leger et al. 2024). However, efforts for systematic integration of the various sources of evidence for human traits have been lacking. Unifying the genetic evidence is particularly relevant to advance our understanding of the ∼7,000 rare diseases due to their largely genetic etiology (80% (Marwaha et al. 2022)), and the fact that unraveling their disease mechanisms has remained challenging in part due to a of lack of patients, unavailable experimental models, and difficulties in linking genotypes to complex phenotypes.

Here, we sought to integrate genetic evidence to unravel mechanisms and prioritize candidate genes for rare diseases. We use representation learning on a protein interaction network coupled with a variational autoencoder to learn latent representations of human traits and other terms. To map the landscape of human traits and disease, we systematically integrate evidence for 11,765 human traits and diseases from various sources (GWAS, Burden, ClinVar and Orphanet), together with 16,009 Gene Ontology (GO) terms, 10,301 mouse phenotypes and 3,575 drugs. We find that similar traits share their latent neighborhoods, revealing an integrated view of human disease in the context of cell biology. Different sources of evidence implicate both shared and specific cell biology for its role in disease. The latent neighbors recover known disease genes across sources of evidence, and we demonstrate how neighborhoods can prioritize candidate genes for rare disorders. Finally, through examples for diabetes, sensorineural hearing loss, and familial long QT syndrome, we illustrate how our methodology can be applied to develop hypotheses for disease mechanisms and to propose disease-relevant experimental models, measurements, and active molecules that may improve diagnosis and treatments.

## RESULTS

As a starting point, we quantified the differences in trait-linked variants and genes for different sources of evidence. As expected, we found that GWAS and ClinVar shared few trait-linked variants for each trait (average Jaccard index = 0.005; Open Targets (OTAR) genetics portal), with ∼96% of GWAS variants being non-coding (intergenic or splicing/intron variants) and ∼81% of ClinVar variants being located in the exome (Fig. 1a). We found a similar discordance between GWAS and ClinVar at the gene level, with the sources of evidence sharing ∼15% of their traits and these traits on average sharing ∼10% of their linked genes (Jaccard index = 0.012) with significant overlaps in linked genes for ∼23% of traits (Fig. 1b - left; one-sided Fisher exact test with BH-adjusted p-values). As expected, we found that traits having genes mostly linked through GWAS were more prevalent than the traits with genes primarily linked through ClinVar (ρ = 0.3, p-value = 1.5e-7; Fig. 1b - right). Expanding the comparison to include other sources of evidence revealed a similar picture - similar sources typically shared traits and their linked genes, while traits and genes largely differ between different sources of evidence (Fig. 1c). Finally, we found that the gene- and protein-level properties of confidently-linked genes differed substantially between sources (Fig. 1d). For example, we found that GWAS linked proteins were over 6x less abundant compared to ClinVar linked proteins (p-value = 4.4e-77; two-sided Welch’s t-test; evidence scores >= 0.5, ignoring proteins linked through both sources), had over 2x less interactions (p-value = 3e-163), with the genes having over 4% lower GC content (p-value = 4.4e-48) and being less essential (p-value = 8.4e-47). Together, these observations demonstrate the discordance of causal genetic evidence for human traits, with largely disjoint sets of linked genes and varying gene characteristics.

**Figure 1.**
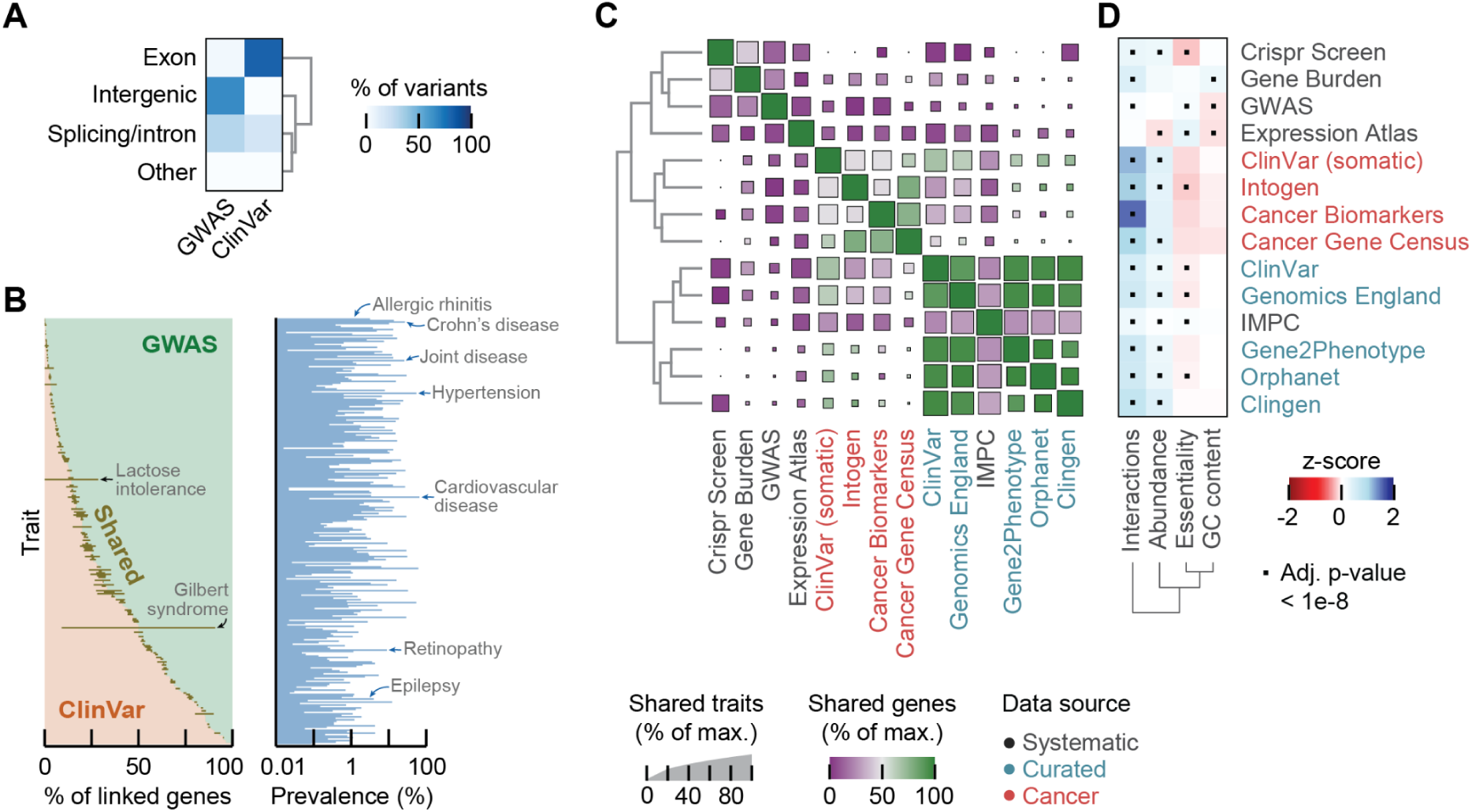
Sources of evidence for human traits differ at the variant and gene-levels. **(A)** Functional consequences of variants linked to traits through ClinVar or GWAS (OTAR). Heatmap shows the percentage of variants categorized as intergenic, exome, splicing/intron, or other. **(B)** Percentage of trait-linked genes that are specific for GWAS (green), ClinVar (orange), or shared (black), together with estimated trait prevalence (FinnGen). Shown are traits reported for both sources of evidence and in FinnGen. **(C)** Percentage of traits shared between sources of evidence (size of squares) and for the shared traits the average percentage of linked genes shared between sources (color). Data sources are annotated as systematic (gray), curated (blue) or cancer-specific (red). **(D)** Characteristics of genes confidently-linked to traits for the sources of evidence. Shown are the number of protein interactions (STRING score >= 0.4), protein abundance (PaxDB whole-organism), gene essentiality (DepMap; average across cell lines), and GC content (Ensembl), z-scored across genes. Interactions and abundances were log-transformed. Black dots show enrichment compared to all other genes not linked to traits for the source of evidence (BH-adjusted p-values < 1e-8; two-sided MWU-test).

### Latent representation of network embeddings for human traits and terms

We sought to further explore the genetic evidence and unravel disease mechanisms for rare diseases by mapping the landscape of human traits. To do so, we used genes confidently associated with traits through evidence from systematic studies (GWAS, burden) and curated datasets (ClinVar, Orphanet), together with genes linked to terms (biological processes, cellular components and molecular function; Gene Ontology) and phenotypes via mouse knockout studies (IMPC) (Methods). In total, we integrated evidence for 11,765 human traits, 16,009 GO terms and 10,301 mouse phenotypes from these 6 sources of evidence. We then created a low-dimensional representation of the traits and terms by coupling a variational autoencoder (VAE) to network embeddings of traits and terms obtained from graph representation learning on a protein interaction network, adjusting for network biases through a re-wiring approach (Fig. 2A). In short, we sampled random walks on a protein interaction network and trained a skip-gram neural network model to obtain 128-dimensional network embeddings of genes (node2vec (Grover and Leskovec 2016)). For each trait and term, we used these network embeddings to score the probabilities of genes to be close to at least one confidently-linked gene, adjusting for network biases by log-normalizing with probabilities analogously derived from 100 randomized networks that were rewired through the configuration model. Finally, we trained a VAE on these normalized network scores to obtain a 128-dimensional latent representation for all human traits and other terms (Methods; Supplementary Figs. 1-2).

**Figure 2.**
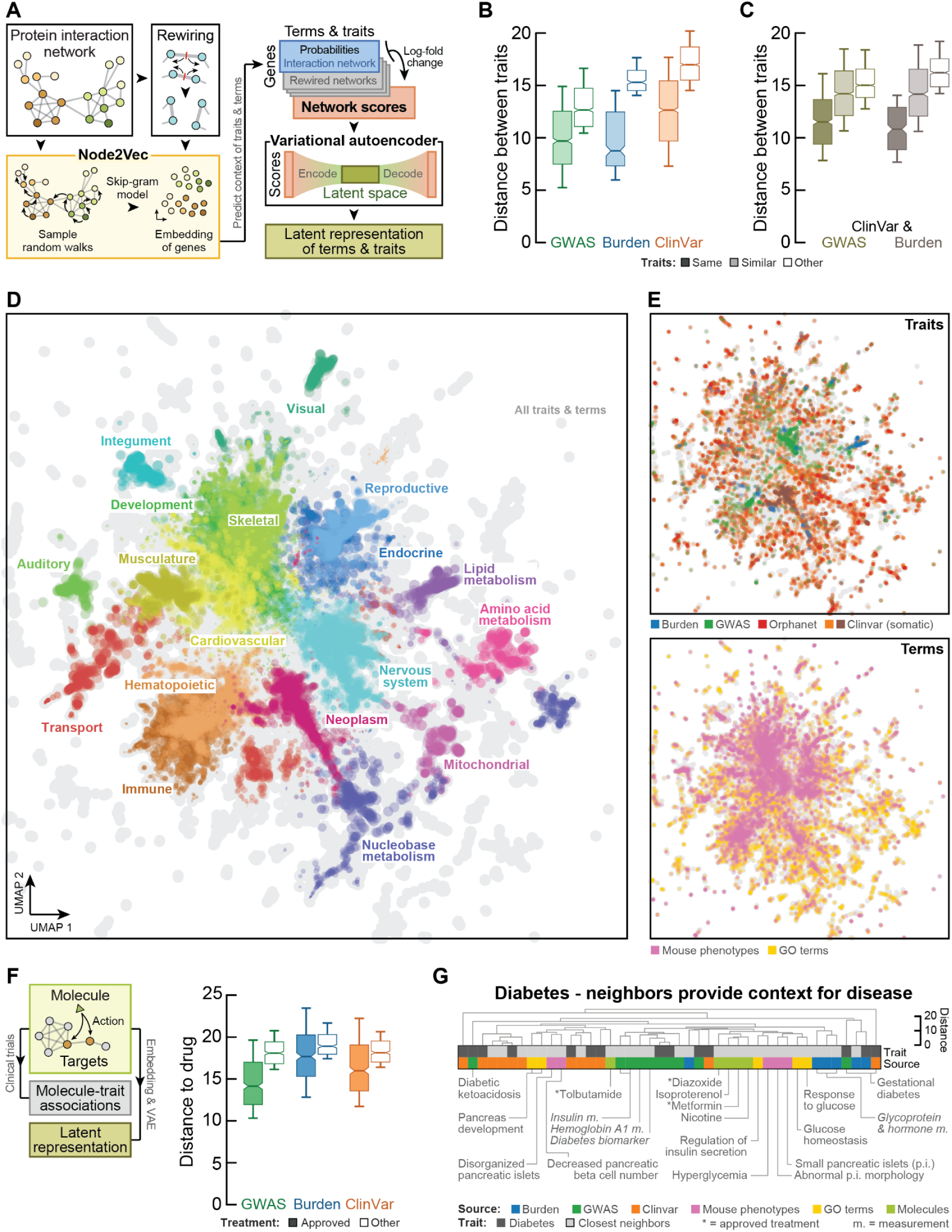
Latent representation of network embedding maps the landscape of human traits in the context of cell biology. **(A)** Schematic of approach. A 128-dimensional network embedding of genes was obtained by training a skip-gram model on random walks sampled from a protein interaction network (node2vec; STRING, IntAct, Reactome, Signor), and analogously for networks re-wired through the configuration model. Network embeddings were used to score the probability of genes to be close to at least one confidently-linked gene for each trait and term. Network scores were computed as the log-fold change of probabilities for the protein interaction network and the re-wired networks, and then used to obtain a 128-dimensional latent representation of traits and terms by training a VAE (Methods). **(B-C)** Distances between traits within sources of evidence for GWAS (green), Burden (blue) and ClinVar (orange) (B) and across sources of evidence for ClinVar-GWAS (dark green) and ClinVar-Burden (brown) (C). Boxplots show average distance for the same trait (dark boxes), between similar traits (light boxes - siblings in the ontology tree) and between other traits (white boxes - traits that are neither siblings or parent-children). Shown are traits having at least 5 confidently-linked proteins and at least one sibling in the ontology tree. **(D)** UMAP of latent representations for traits and terms. Traits and terms were colored according to their most likely broad category from the respective ontology trees. Probabilities were estimated from a gaussian kernel density for each of the categories. Colored dots were sized proportionally to the probability of the respective category, showing only dots having probabilities exceeding the median probability of all traits and terms (Methods). Background shows all traits and terms (grey). Margins of UMAP were trimmed for visualization to show all systematically annotated traits plus one unit distance (97.6% of all traits and terms - Methods). **(E)** UMAP of latent representations as in (D), annotating sources of evidence for traits (top) and terms (bottom). Shown are traits from GWAS (green), Burden (blue), ClinVar (somatic) (orange and brown respectively), Orphanet (red), mouse phenotypes (pink) and GO terms (yellow). Background shows all traits and terms (grey). **(F)** Diagram shows schematic of approach. Network embedding and VAE from (A) were re-used to create latent representations of molecules through their confidently-linked protein targets (STITCH score >= 700). Boxplots show average distance of drugs to traits they were approved for (Clinical Stage >= 4; OTAR) compared to other traits, for GWAS (green), Burden (blue) and ClinVar (orange). Shown are drugs having at least 5 confidently linked targets. **(G)** Exemplary application of the proposed methodology. Shown are traits annotated as diabetes mellitus (EFO_x - dark grey), together with the 5 nearest neighbors of diabetes for each source of evidence (light grey). Metric was the Euclidean distance in latent space to the median of traits annotated as diabetes mellitus (EFO_0000400). Traits and terms were clustered through complete-linkage clustering of latent representations. Shown are traits from GWAS (green), Burden (blue), ClinVar (orange), mouse knockout phenotypes (pink), GO terms (yellow) and molecules (light green). Biomarkers and measurements shown in italics, approved treatments annotated with asterisks. Boxplots in (B,C,F) show 10-th and 90-th percentile (whiskers) with the median and s.e. (notches). Metric is the Euclidean distance of latent representations.

We found that similar traits - traits sharing a parent in the ontology graphs (Methods) - were mapped closer together in the latent space compared to unrelated traits for GWAS (1.6-fold; p-value = 6.9e-181; one-sided paired t-test), Burden (1.8-fold; p-value = 3.6e-88) and ClinVar (1.6-fold; p-value = 0) (Fig. 2B). Moreover, we found similar results when comparing distances of the same trait across different sources of evidence. Indeed, traits with evidence from ClinVar were typically closer to the same trait with evidence from GWAS (1.3-fold; p-value = 1.3e-130) and Burden (1.5-fold; p-value = 1e-75) compared to the other traits from the same sources respectively (Fig. 2C). This extended beyond simple overlaps in disease-linked genes (median jaccard index = 0.01 for the 21% of pairs of traits sharing any genes), and regressing these overlaps out of the distances still mapped similar traits closer than unrelated traits by ∼27% between the common and rare variant studies (Supplementary Fig. 3). Together, these observations suggest that similar traits share the same neighborhood in the latent space, despite the sources of evidence diverging at the variant and gene level. Furthermore, we confirmed that the relative distances between similar and unrelated traits from our network scores and low-dimensional representations significantly outperformed the distances from traditional approaches such as network propagation (e.g., improved the distances for the same trait relative to other traits across sources of evidence for ClinVar and GWAS by ∼12%; p-value < 2.4e-55 - Supplementary Figs. 4-5).

### Mapping the landscape of human traits and terms

Having established that the latent representation of network scores for similar traits share the same neighborhood, we annotated the human traits, mouse phenotypes and GO terms with their broad categories from the respective ontology graphs and visualised them through a UMAP (Fig. 2D; Supplementary Tables 1-2 - Methods). This visualization revealed a striking landscape of human traits and terms, with closely related organ systems sharing their locations, such as for the immune and hematopoietic systems, reproductive and endocrine systems, or the musculature and cardiovascular system. Similarly, we found that the skeletal system shared its location with development, while neoplasms were next to the metabolism of nucleobases. The same map revealed that the human traits intermixed with mouse phenotypes and GO terms (Fig. 2E). We found similar maps when preserving relative distances through a PCA of the network scores - the first principal component separates most pairs of broad categories of traits and terms, with related categories having more similar scores compared to unrelated categories (Supplementary Figs. 6-7).

Next, we hypothesized that we could use the pre-trained network embeddings and VAE to map additional sources onto the same landscape. To test this, we took the molecule-protein associations (molecules acting on proteins from STITCH; scores >= 700 (Szklarczyk et al. 2016)), and created network scores and latent representations for the molecules through the VAE that we trained on the traits and terms. In the latent space, we found that drugs were mapped closer to the traits for which they were approved compared to other traits that had approved treatments (Fig. 2F - drug-trait associations from OTAR; Clinical Stage >= 4), especially for traits from GWAS (24% closer; p-value = 1.4e-56; one-sided MWU-test) and ClinVar (13% closer; p-value = 1.4e-22). For example, compared to other diseases and for both the ClinVar and GWAS, we found that Rosuvastatin mapped close to hyperlipidemia (among the top 2%), while Propafenone mapped close to cardiac arrhythmia (top 1%), Vildagliptin mapped closer to type 2 diabetes compared to the more commonly used Metformin (top 1% and 7% respectively), and Nifedipine and Hydralazine mapped closer to hypertension through ClinVar (top 4%) compared to GWAS (>6%). Together, these observations demonstrate that the pre-trained network embedding and VAE map unseen evidence into the appropriate neighborhood of the existing samples, and may be applied to include additional traits, terms and likely other ontologies.

### Latent neighbors provide context for disease

As an example of applying our methodology for studying disease, we selected the 14 traits that had latent representations and were annotated as diabetes mellitus (children of EFO:0000400). Besides type I, type II, gestational and general diabetes for both GWAS and ClinVar, this also included insulin-resistant, neonatal insulin-dependent, monogenic and rare genetic diabetes (ClinVar), diabetic ketoacidosis (GWAS), and general diabetes mellitus (Burden). For all other traits and terms, we computed the euclidean distance of the latent representations to these diabetes traits. Taking the median of distances to the diabetes traits, we then selected the 5 traits or terms for each source of evidence whose latent representation was nearest to diabetes. The selected traits and terms typically shared few of the confidently-linked genes with the diabetes traits (jaccard index ∼0.04 p/m 0.01), and 59% of the selected traits and terms would not have been selected based on their jaccard index or enrichment with the confident diabetes genes (Supplementary Table 3). However, all of the selected neighbors were highly relevant for diabetes and its disease mechanism (Fig. 2G). From the GO terms, we found processes related to pancreatic development, regulation of insulin secretion, and the response to glucose and its homeostasis. The nearest neighbors also proposed mouse models having phenotypes related to disorganized and abnormal morphology of pancreatic islets, hyperglycemia, and a decreased pancreatic beta cell number. From GWAS and Burden tests, we identified biomarkers and several measurements known to be relevant for diagnosing diabetes, such as hemoglobin A1 (Gilstrap et al. 2019), glycoprotein and insulin measurements. Finally, we found that three of the five nearest molecules were approved drugs (Diazoxide (for hypoglycemia), and Tolbutamide and Metformin (for type II diabetes) (Zdrazil et al. 2024)), while the other two molecules are known risk factors for developing insulin resistance (isoproterenol and nicotine (Hoff and Koh 2018; Chen et al. 2023)). Together, these neighbors provide context for diabetes and its disease mechanism by identifying related cellular processes, experimental models, diagnostic measurements, and active molecules that are treatments or risk factors. Overall, this example illustrates how our methodology can be applied to develop hypotheses for disease mechanisms and may propose disease-relevant experimental models, diagnostics, and active molecules that are not found through simple overlaps in gene sets.

### Related traits converge onto the same modules of functionally related genes

Having established that the network scores and latent representations of traits and terms can be used to integrate sources of evidence, we hypothesized that the different sources of evidence for a trait - due to their discordance - may highlight different aspects of the same cell biology (Barrio-Hernandez et al. 2023; Leger et al. 2024). To test this, we clustered the protein interaction network into functional protein modules and linked the modules to traits and terms through the network scores and gene-set enrichment analysis (GSEA) (Fig. 3A; Supplementary Tables 4-5 - Methods). Indeed, we found that closer traits shared more protein modules (Pearson ρ = -0.59; p-value = 0.0), and that similar traits shared more modules compared to unrelated traits for the same source of evidence (Fig. 3B), such as for ClinVar (7.3-fold difference of Jaccard index on average; p-value = 3.2e-264 - one-sided paired t-test) and GWAS (4.3-fold; p-value = 1.6e-133). We found the same trend for the same and similar traits compared to unrelated traits across sources of evidence (Fig. 3C), such as for GWAS and Clinvar (2.5-fold and 1.6-fold for difference for the same and similar traits respectively; p-values = 1.9e-86 and 6.4e-89). These observations demonstrate that similar traits share substantially more functional protein modules compared to unrelated traits. Moreover, we found that these differences across the sources of evidence are not driven by overlaps in confidently-linked genes or the similarities of the network scores (Supplementary Fig. 8), suggesting that the shared modules may be linked to the traits through different aspects of their biology.

**Figure 3.**
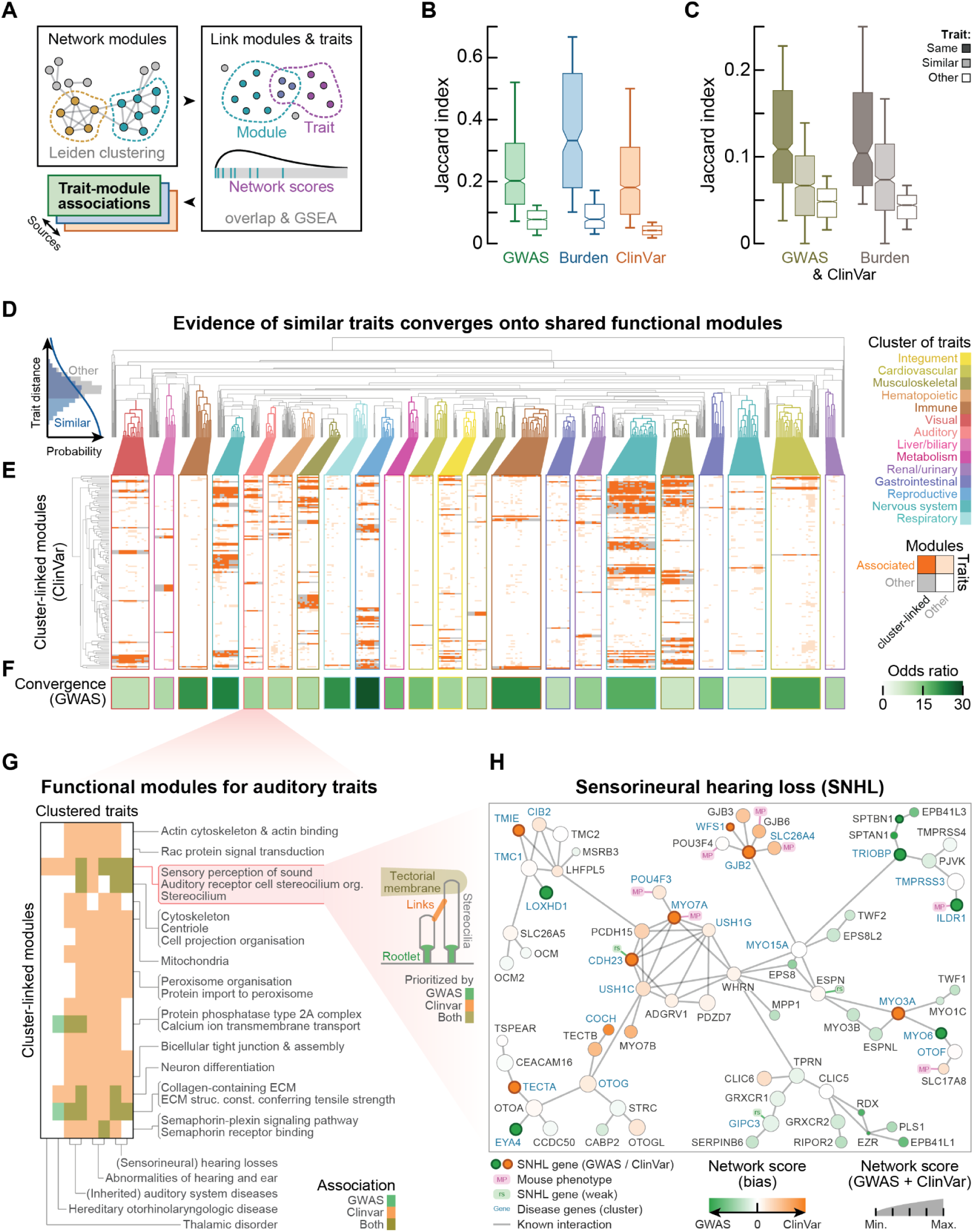
Related traits converge onto the same functional protein modules. **(A)** Schematic of approach. The protein interaction network is clustered into functional gene modules. Traits and terms are linked to modules through shared genes and gene-set enrichment analysis with the network scores (Methods). **(B-C)** Shared modules between traits within sources of evidence for GWAS (green), Burden (blue) and ClinVar (orange) (B), and across sources of evidence for ClinVar-GWAS (dark green) and ClinVar-Burden (brown) (C). Shown is the average jaccard index of modules linked to the same trait (dark boxes), similar traits (light boxes - c.f. Fig. 2B) and between other traits (white boxes). Shown are traits having at least 5 confidently linked proteins and with at least one sibling. Boxplots show 10-th and 90-th percentile (whiskers) with the median and s.e. (notches). **(D)** Probability of traits to be similar as a function of distance (left) and dendrogram clustering 570 ClinVar traits also reported for GWAS (right; excluding neoplasms). (left) Histogram shows average distance to similar traits (blue; sharing a parent in the ontology tree) compared with average distance to other traits (grey; traits that are neither siblings nor parent-children). Probabilities are from a logistic model fitted to the distances (solid blue line). (right) Selected clusters contain at least 8 traits that are likely similar (probability > 0.5) and colored according to strongest enrichment of traits from a broad category from the ontology tree (c.f. Fig. 2D). Traits were clustered through complete-linkage clustering. Metric is the Euclidean distance of latent representations from ClinVar. **(E)** Associations of cluster-linked modules. Modules are linked to clusters (grey) when enriched with associations to traits of a cluster (dark orange). Other trait-module associations are shown in light orange. Modules are clustered through complete-linkage clustering of the associations using the Manhattan distance. **(F)** Enrichment of associated functional modules between GWAS and ClinVar for traits in each subcluster. Shown is the odds-ratio (one-sided Fisher exact test; all BH-adjusted p-values < 5e-5). **(G-H)** Example for convergence of GWAS and ClinVar evidence onto a functional module (stereocilia with Sensorineural Hearing Loss; SNHL). (G) Heatmap shows associations between clustered auditory traits (c.f. Fig. D) and cluster-linked modules (c.f. Fig. E). Associations are through evidence from GWAS (green), ClinVar (orange) or both (olive). Protein modules were named through enrichment of GO terms (Methods). Cartoon illustrates stereocilial substructures significantly prioritized by the respective sources of evidence. (H) Network of protein interactions for proteins in the stereocilium module (G - Methods). Nodes are colored to the bias of network scores for being higher for ClinVar (light orange) or GWAS (light green), sizes are proportional to average network score from ClinVar and GWAS. SNHL disease genes are colored for ClinVar (dark orange) and GWAS (dark green). Nodes are annotated for having weak SNHL variants (‘rs’), associations to SNHL in mice (‘MP’) or being confidently-linked to at least 5 other auditory traits in the cluster (blue text).

To further explore the convergence of traits onto shared functional modules of proteins, we fitted a logistic model to the latent distances between ClinVar traits to score the likelihood for any pair of ClinVar traits to be similar or unrelated (Fig. 3D - left histogram; Methods). We then clustered the 570 ClinVar traits that also had evidence from GWAS and that were not neoplasms (Fig. 3D - right dendrogram). By combining the model and clustered traits, this dendrogram revealed subclusters of traits that were likely to be similar and that were enriched for common annotations in the ontology graph (Fig. 3D - colored subclusters; Methods), such as several subclusters enriched for traits related to the immune, nervous and cardiovascular systems. For such subclusters, we found gene modules enriched with associations to the clustered traits (Fig. 3E; Supplementary Tables 6-7; Methods). Some of these cluster-linked modules were general, while others were specific to clusters of particular types of traits. Labeling modules through their enrichment with genes from GO terms (Methods), we found for example modules for DNA repair, translation and respiration to be fairly unspecifically linked, while modules such as for bile acid secretion, keratinization, ossification, and GABA receptor activity were more specifically linked to clusters of the liver/biliary system, the integument, musculoskeletal and nervous systems respectively (Supplementary Fig. 9).

Next, we tested whether the subclusters of traits converged onto the same functional modules through evidence from GWAS or ClinVar. Indeed, we found that each subcluster of traits shared their associated functional modules between the sources of evidence (average odds-ratio 14.7 p/m 1.2; BH-adjusted p-values < 1.4e-03 across all subclusters - one-sided Fisher exact tests; Methods) (Fig. 3F). For example, we found that the evidence for reproductive disorders converged most strongly onto the same functional modules (odds-ratio 28.2), similar to subclusters of disorders of the immune system (odds-ratios 22.0 and 21.1). Strikingly, evidence for subclusters of disorders of the nervous system such as epilepsies (22.8) and neurological disorders (18.2) converged more strongly onto the same functional modules compared to other diseases of the nervous system such as movement disorders (6.0) and diseases of the musculoskeletal system (7.6, 10.5, and 11.4). Together, these observations corroborate the findings from previous studies exploring the pleiotropy of human cell biology through functional protein modules (Barrio-Hernandez et al. 2023). Moreover, these analyses show that traits with evidence from common and rare variant studies converge onto functional protein modules despite having small overlaps in confidently-linked genes. Finally, looking at the same traits with evidence for GWAS, Burden and ClinVar, we found that the ClinVar and Burden evidence (median odds-ratio = 50.9; Fisher exact tests) converged more strongly onto shared functional modules compared to the Burden and GWAS (19.8; p-value = 3.6e-3) or ClinVar and GWAS evidence (15.6; p-value = 7.0e-8 - two-sided MWU test), suggesting that rare variant analysis of population studies may in part bridge the differences in trait-linked genes identified through GWAS and clinical genetic testing (ClinVar).

### Converging evidence for sensorineural hearing loss (SNHL)

To further explore the convergence of the common and rare variant studies, we looked at aspects of cell biology prioritized by the sources of evidence. Specifically, we took the subcluster of hearing disorders and focussed on sensorineural hearing loss (SNHL) with a protein module for stereocilia that was linked to the cluster through both GWAS and ClinVar (Fig. 3G - heatmap). The GWAS and ClinVar evidence for SNHL does not share any of the confidently-linked genes (56 genes in total). SNHL accounts for roughly 90% of reported cases of hearing loss, affecting roughly 5% of the world population (Sheffield and Smith 2019). Unraveling the genetics of SNHL is of clinical importance because SNHL has a genetic etiology in over 50% of cases and is highly heterogenic, with limitations of (genetic) screening methods causing cases and at-risk carriers to remain undetected (Smith et al. 2005; Linden Phillips et al. 2013; Koffler et al. 2015; Sheffield and Smith 2019). The functional module for stereocilia is appropriate in the context of SNHL, as stereocilia are responsible for sound transduction by acting as mechanosensors on the surface of hair cells (McGrath et al. 2017). As we will demonstrate, the network scores for SNHL revealed stereocilial substructures prioritized by both GWAS and ClinVar (tectorial membrane), but also source-specific prioritization of components such as stereocilia roots (GWAS) or tip links (ClinVar) (Fig. 3G - cartoon).

We looked for common and source-specific perspectives on SNHL by the common and rare variant studies. To do so, we visualized the stereocilium module as a network using known protein-protein interactions (Fig. 3H - Methods; OTAR confidence score > 0.95), and annotated the proteins using the network scores for SNHL to show their average score (size) and score bias for GWAS (green) or ClinVar (orange) (Fig. 3H - Methods). We found that the proteins in the stereocilium module were highly ranked for SNHL through both GWAS and ClinVar (p-value = 8.6e-49 and 2.6e-48 respectively; one-sided MWU-tests ignoring confidently-linked proteins), with 24 proteins having supporting evidence of at least 5 of the other traits in the cluster of hearing disorders (Fig. 3H - blue gene names; Supplementary Table 8).

The visualized network revealed functional subgroups of genes prioritized by both sources of evidence or more specifically by GWAS or ClinVar. For example, after excluding the confident SNHL disease genes, we found functional subgroups prioritized specifically by ClinVar, such as highly interconnected genes involved in hair-bundle links and the mechanoelectrical transduction (MET) channel (p-value = 7.6e-3; two-sided paired t-test), or connexin and chloride transport (p-value = 6.6e-3). In contrast, functional subgroups scoring higher for GWAS involved genes localized at stereocilium rootlets (p-value = 1.3e-2 - PJVK, RIPOR2, CLIC5, GRXCR2, TPRN, RDX) (Park and Bird 2023; Michalski and Petit 2015) or actin-interacting proteins (p-value = 3.9e-3). Finally, as an example of functional subgroups prioritized by both GWAS and ClinVar, we found that proteins of the tectorial membrane ranked higher than other proteins in the stereocilium module (p-value = 1.9e-04; one-sided MWU-test) (Michalski and Petit 2015; Bian et al. 2025), with OTOG, OTOA and CEACAM16 ranking in the top 5 for both sources of evidence. Thus, the tectorial membrane is prioritized as functionally important for SNHL through both GWAS and ClinVar, without substantial differences between these sources (p-value = 0.11; two-sided paired t-test). The tectorial membrane - a gel-like ECM attached to stereocilia - is essential for hearing by stimulating hair cells and facilitating sound transduction. Indeed, disruptions of the tectorial membrane are tightly linked to hearing losses, opening avenues for potentially novel biomarkers and gene therapies (Bian et al. 2025; Michalski and Petit 2015). Overall, these examples illustrate how common and rare variant studies converge onto shared functional modules of cell biology and highlight specific aspects of its cell biology.

### Latent neighbors prioritize disease-linked genes

With the close latent neighbors providing context for disease from the perspective of the different sources of evidence, we hypothesized that we could use the neighbors to prioritize candidate disease genes for (rare) disorders. To do so, we first tested whether the network scores of genes for traits could recover known disease-linked genes. For each source of evidence we selected traits having at least 20 confidently-linked genes and created n=100 sub-samples for each trait by withholding subsets of 25% of the trait-linked genes and computing network scores using the remaining linked genes (Supplementary Fig. 10; Methods). We found that the network scores of sub-samples could recover the withheld trait-linked genes, and this recovery was trait-specific as they outperformed the recovery of other frequent disease genes for the same source of evidence (Methods). Indeed, we found that the network scores recovered confidently linked genes in a trait-specific manner for GWAS (average AUC = 0.77 and z-score = 6.19), Burden (0.84 and 4.57) and ClinVar (0.85 and 5.87), but also genes that were less confidently linked to the traits, in particular for the rare diseases (0.73 and 3.91 - Fig. 4A). Finally, we confirmed that the network embedding scores outperformed the scores derived from network propagation for recovering trait-linked genes, with the network scores substantially improving the average precision-recall (2.2-fold; p-value = 2.2e-20; Supplementary Figs. 10-11). Together, these results suggest that the network scores can be used to prioritize disease genes in a trait-specific manner.

**Figure 4.**
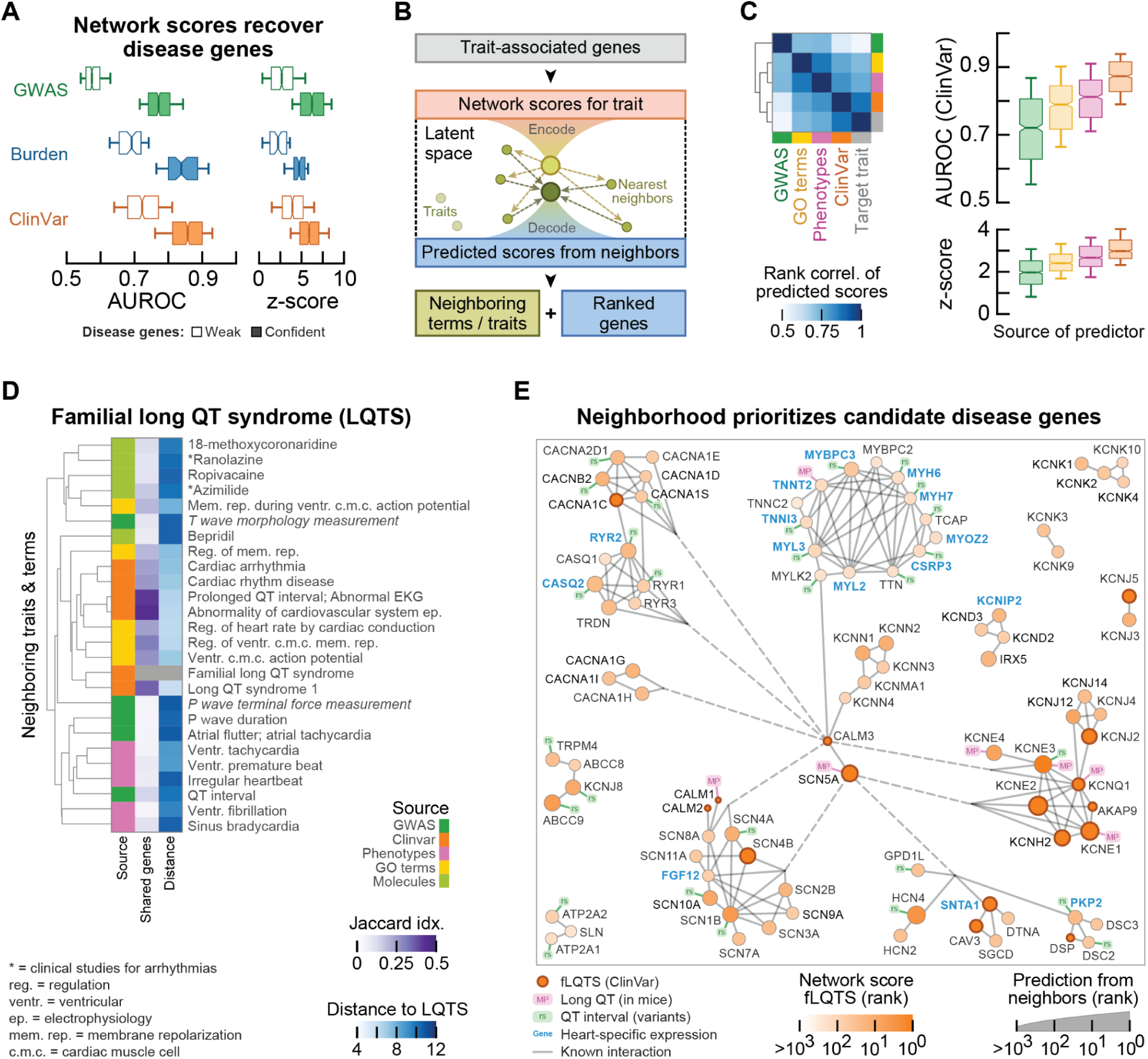
Close neighbors prioritize candidate disease genes. **(A)** Recovery of trait-linked genes using network scores for GWAS (green), burden (blue) and ClinVar (orange) traits. Shown is the recovery of withheld confidently-(colored; scores >= 0.5) and weakly- (white; scores < 0.5) linked genes (AUROC) by the network scores, compared with the recovery for genes sampled from their frequencies for the respective sources of evidence (z-score). **(B)** Schematic of approach. Trait-linked genes are used to compute network scores and latent representation for a trait. For a given source of evidence, the 5 nearest neighbors are selected and averaged in latent space. The neighbors are averaged and decoded to obtain predicted network scores for the trait. **(C)** Heatmap (left) shows rank correlation of network scores for a ClinVar trait with the predicted scores from the nearest neighbors of its sub-samples, for neighbors of the various sources of evidence. Boxplots (right) show recovery (AUROC) of withheld linked genes by the predicted scores from nearest neighbors, compared with the recovery by distant neighbors (z-score). Neighbors are from GWAS (green), ClinVar (orange), GO terms (yellow) and mouse phenotypes (pink). **(D-E)** Example of prioritizing candidate disease genes for a rare disorder (familial long QT syndrome; fLQTS). (D) Dendrogram shows close neighbors selected from GWAS (green), ClinVar (orange), GO terms (yellow), mouse phenotypes (pink), and molecules (light green). Neighbors were clustered through complete-linkage clustering with the latent representations and the Euclidean distance. Heatmap shows distances of neighbors to fLQTS (blue) and the confidently-linked genes shared between the nearest neighbors and fLQTS (jaccard index - purple). **(E)** Network of confident protein interactions (confidence score >0.95 - grey edges) for highly ranked proteins (among top 100) according to the predicted network scores for at least two sources of evidence. Nodes are colored according to the rank of the fLQTS network score (ClinVar) and sized proportional to the rank of the average predicted scores from the nearest neighbors. Nodes are annotated as fLQTS disease genes (brown edge), having other variants linked to the QT interval (‘rs’), and with associations to long QT in mice (‘MP’). Genes highly and specifically expressed in heart muscle are shown in blue. Boxplots in (A,C) show 10-th and 90-th percentile (whiskers) with the median and s.e. (notches). Metric is the Euclidean distance of latent representations.

Having confirmed that the network scores recover both confident and weak trait-linked genes (Fig. 4A), that similar traits are closer than unrelated traits (Fig. 2) and that the sources of evidence highlight different trait-relevant aspects of cell biology (Fig. 3), we next sought to prioritize candidate disease genes for rare diseases using predictions from network scores of similar traits and across sources of evidence. To do so, we used trait-linked genes to compute the latent representation of a trait and selected its 5 nearest neighbors in the latent space. We then averaged the latent representations of these neighbors, which we then decoded with the VAE to obtain predicted network scores for a trait based on the evidence from similar traits and other sources of evidence (Fig. 4B). To test the proposed methodology, we used the latent representations of sub-samples for the ClinVar traits to select the nearest latent neighbors for each sub-sample. We then used these nearest neighbors to predict network scores and recover the known trait-linked genes that were withheld for the sub-sample (following Fig. 4A; Methods).

We found that the network scores of ClinVar traits were recapitulated by the scores predicted from the nearest neighbors of its sub-samples, in particular when using neighboring ClinVar traits (median Spearman correlation ρ = 0.87) (Fig. 4C - left). Moreover, these predicted scores from the nearest neighbors could recover the unseen disease-linked genes, and this recovery was specific to the nearest neighbors compared to more distant traits (Fig. 4C - right). Indeed, we found that the withheld trait-linked genes were recovered by the predicted scores from the nearest neighbors of GWAS (average AUROC = 0.71, z-score 1.97), GO terms (0.78 and 2.45), mouse phenotypes (0.81 and 2.71), and other ClinVar traits (0.87 and 3.08). As expected, the recovery of linked genes worsened when using increasingly distant traits, and the nearest ClinVar neighbors significantly outperformed the other sources of evidence at the same distances (Supplementary Fig. 12). Moreover, we confirmed that the nearest neighbors prioritized genes for rare diseases beyond simple overlaps in linked genes (e.g., AUROC = 0.84 after regressing out the overlap in linked genes with nearest ClinVar neighbors; Supplementary Figs. 13-15). Furthermore, combining neighbors from different sources of evidence did not further improve the recovery of trait-linked genes (p-value = 0.55; two-sided Welch’s t-test - Supplementary Fig. 15), likely due to the sources of evidence highlighting different aspects of cell biology (Fig. 3). Indeed, compared to ClinVar, GWAS evidence may be more likely to prioritize genes whose variants are risk factors or modulate the phenotype for the rare diseases. Finally, we found that decoding the latent representations of nearest neighbors outperformed other approaches such as averaging the network embedding scores or network propagation scores of nearest neighbors (Supplementary Fig. 16). Together, these analyses show that the nearest latent neighbors of rare diseases prioritize disease genes in a trait-specific manner and across sources of evidence.

### Prioritizing candidate genes for familial Long QT Syndrome

To demonstrate how the proposed methodology can be applied for prioritizing candidate disease genes for rare disorders, we looked at familial Long QT Syndrome (familial LQTS). LQTS is a rare heritable condition characterized by dysfunctioning cardiac repolarization (Baskar and Aziz 2015; Asatryan et al. 2018), leading to abnormal QT interval length and with high risks of developing heart arrhythmia and cardiac arrest (Galić et al. 2021; Tan et al. 2025). The prioritization of novel candidate genes is of clinical importance as approximately 25% of cases lack a genetic diagnosis (MacRae 2009; Wilde et al. 2022). As before, we used its confident disease genes from ClinVar to compute network scores and its latent representation, from which we then selected the 5 nearest neighboring traits and terms for each source of evidence (Fig. 4D). We found all traits and terms to be closely related to cardiac function and electrophysiology, but generally shared few genes with fLQTS. Among others, we found four processes related to membrane repolarization and action potential of cardiac muscle cells (GO terms; Jaccard index 0.24), five heart rate phenotypes (mouse phenotypes - 0.07), four characteristics of electrocardiograms including two diagnostic measurements (GWAS - 0.08) and four other cardiac arrhythmias (ClinVar - 0.34). Finally, we found two drugs approved for cardiac arrhythmias (Ranolazine and Azimilide (Zdrazil et al. 2024)).

As before, we averaged these nearest neighbors in latent space and decoded them using our VAE to obtain predicted scores for fLQTS for each source of evidence. The predicted scores strongly correlated with the network scores of fLQTS (Spearman ρ = 0.95) and recovered the confident fLQTS-linked genes (AUROC = 0.95; average PR = 0.35). To prioritize candidate disease genes, we selected the genes that were among the top 100 highest ranking genes according to the predicted scores for at least two sources of evidence. We then visualized these genes through a network of known protein-protein interactions (Fig. 3E; Supplementary Table 9 - Methods), and annotated the proteins using their ranks for fLQTS (color) and the predicted ranks from the decoded neighborhoods (size).

The visualized network contains several clusters of highly interconnected genes, in particular for voltage gated channels of calcium, sodium and potassium and for the troponin complex and cardiac myosin. After removing the n=16 confident disease genes from fLQTS, the visualized network is enriched for genes highly expressed (among top 5%) and specific (z-score > 2) to the heart muscle (odds-ratio = 10.9, p-value = 1.1e-10), and enriched for other genes having variants with supporting evidence linking them to the (long) QT interval (odds-ratio 56.2; p-value = 4.9e-35; one-sided Fisher-exact tests). Indeed, some of the highly ranked genes were confidently associated with long QT but not among the disease genes of fLQTS (e.g., RYR2, MYBPC3 and MYH6 - prioritized through the predicted scores), and 23 other genes in the network had weaker evidence supporting them as causal for long QT traits. Given their interactions with other fLQTS-linked genes, these could be prioritized as more likely to be causal due to their functional roles. For example, we found that KCNE3 had several variants with weaker evidence for LQTS. KCNE3 has high confidence interactions with 5 known LQTS-linked genes and has been associated with long QT through knockout studies in mice (Groza et al. 2023). Indeed, KCNE3 was the highest ranked gene for the network scores for fLQTS and was ranked 4th according to the predicted network scores from the other sources of evidence. We also found other cases with (f)LQTS reported in literature that lacked genetic diagnosis and are currently not linked to LQTS in Open Targets or ClinVar, but had mutations in highly ranked genes, such as for SCN10A (Abou Ziki et al. 2018), KCNJ8 (linked to sudden infant death (Tester et al. 2011)), compound mutations for ABCC9 and KCNJ8 (ranked 4th and 8th respectively) (Hu et al. 2014), or variants in HCN4 (ranked 2nd) acting as modifiers of the LQTS phenotype in conjunction with mutations for known fLQTS genes KCNH2 or SCN5A (i.e., carriers are more strongly affected) (Copier et al. 2025; Bukaeva et al. 2025). Together, these results suggest that screening for variants in the highly ranked genes may be of clinical importance for diagnosis and treatment for LQTS patients.

### Systematic prioritization of candidate genes for rare diseases

Finally, following the same approach as for fLQTS, we used the other sources of evidence to systematically predict latent neighborhoods of traits and terms for n=477 common and rare diseases, which we then used to prioritize candidate genes (Supplementary Table 10 - Methods). This analysis compiled 498 gene-disease pairs that are enriched with weaker prior evidence (odds-ratio = 27.5, p-value = 0.0; one-sided Fisher exact test), 225 of which were further supported by functional associations of the protein with other proteins confidently-linked to the disease. For example, through the latent neighborhoods, we prioritized COL4A3 (rs200107989) for Stickler syndrome, SOST (rs76410205) for osteoporosis, MSX1 for Craniosynostosis (rs184700656), SCN9A for Dravet syndrome (e.g., rs200689195), GRIN2A (rs1057519551; rs531782747) and GABRD (rs1057519556) for epileptic encephalopathy, ABCG5 (rs11887534) for familial hypercholesterolemia, APOA5 for familial lipoprotein lipase deficiency (rs1246031494), CR1L (rs679515; rs3818361; rs6701713) for Alzheimer disease, and CFI (rs760801046) for congenital afibrinogenemia. The compilation of prioritized gene-disease pairs with supporting evidence is available through Supplementary Tables 11-12.

Overall, the examples demonstrate how our approach can be applied to propose traits and terms from different sources of evidence that combine into an integrated view of (rare) disease, linking diseases to cell biology, experimental models, active molecules, diagnostic measurements and other traits with supporting evidence from clinical and population-level studies. Moreover, these closely related traits and terms prioritize novel candidate genes that are enriched with disease-relevant variants and may improve diagnosis and treatments.

## DISCUSSION

Previous studies have noted a significant phenotype-specific overlap between mendelian disorder genes and GWAS-linked genes (Freund et al. 2018). In our analysis, we find significant overlaps in linked genes for ∼23% of traits with genes having GWAS and ClinVar information. These observations are limited by the fact that they are not studied within the same cohort. More recent cohort studies based on gene burden analysis have equally observed some degree of overlap with rare-variant associated genes showing significant enrichment for common-variant heritability (Weiner et al. 2023). Despite such observations, the degree of overlap between common and rare variant studies remains very low, in part due to differences in their approach of establishing genetic evidence. It has been recently proposed that rare variant burden tests and GWAS tend to prioritize different sets of genes, in part also due to technical differences in methodologies (Spence et al. 2026). Here, we note as well that different methodologies identify significant overlaps in linked genes for some traits but that the sets of genes are mostly disjoint.

We developed a deep learning method for systematically integrating discordant sources of genetic evidence through latent representations. By coupling a VAE to network embeddings obtained from graph representation learning on a protein-interaction network, we obtain low-dimensional latent representations of human diseases and other traits and terms, and apply these to prioritize candidate genes for rare diseases. After training the model, our approach is dramatically faster for large datasets and outperforms traditional graph-based methods such as network propagation in recovering disease-linked genes. The predictive power of the latent representations is further supported by the embedding of drugs near their indications.

Despite the lack of overlaps in genetic evidence between common and rare variant studies, we demonstrated that similar traits share their latent neighborhoods and that the sources of genetic evidence converge onto functional protein modules. These effects extend beyond simple overlaps in linked genes, suggesting shared cell biology and disease mechanisms for similar traits and across sources of evidence. Indeed, convergence of genetic evidence onto shared mechanisms for similar diseases and across sources is supported by our and other studies (Goh et al. 2007; Menche et al. 2015; Wright et al. 2025). We observed this convergence to be stronger for genes linked through rare variants from population studies (Burden) and clinical genetic testing (ClinVar) compared to genes linked through common variants (GWAS), suggesting that Burden testing may bridge some of the discordance in trait-linked genes identified via clinical genetic testing and population studies.

Our example for hearing loss demonstrates how common and rare variants can map to subtly different aspects of the same processes. Additionally, varying degrees of convergence for different groups of nervous system diseases and stronger convergence for diseases of the immune system compared to the musculoskeletal system may reflect varying degrees of this effect (e.g., differences in genetic heterogeneity and constraints onto modules). These observations may relate to the omnigenic model (Boyle et al. 2017) with traits arising from core genes affected by rare variants (ClinVar) and a plethora of peripheral genes that modulate the phenotype through more common variants (GWAS).

The presented methodology is highly flexible and could be extended with additional evidence and may be improved for specificity. For example, the protein interaction networks are not context-specific, although diseases often manifest in specific cell-types or tissues (Hekselman and Yeger-Lotem 2020). With ongoing efforts for generating context-specific networks (Laman Trip et al. 2025; Skinnider et al. 2021; Huttlin et al. 2021; Schaffer et al. 2025), our methodology can be augmented with such networks to create cell-type or tissue-specific embeddings that are appropriate within the context of disease. Furthermore, having established that similar traits tend to share their latent neighborhoods across sources of evidence, these sources may be more tightly integrated through a contrastive loss that penalizes the distance for similar traits across sources of evidence. Alternatively, through modified architectures or loss functions, the latent representations may be applied for generative disease models or to translate between sources by reconstructing network scores for the same trait from another source of evidence. Finally, by integrating additional modalities such as differential expression data (Hu et al. 2025), the trained model can be extended to generate hypotheses for diagnostic measurements or drug repurposing and combination drugs. These extensions may be further applied in other contexts, for example through representations of an individual’s genetic makeup or blood proteomics to improve diagnosis and advance personalized therapeutics.

## METHODS

### Evidence for associations of genes with traits and terms

Evidence of gene-trait associations, including indirect association, was collected from the Open Targets (OTAR) Genetics Portal (version 24.09; https://platform.opentargets.org/) (Ghoussaini et al. 2021; Mountjoy et al. 2021). These included gene_burden, ot_genetics_portal (GWAS), eva and eva_somatic (ClinVar), and orphanet. The indirect evidence was filtered for confident associations (OTAR score >= 0.5) and further filtered for genes that had network embeddings (Network embedding of proteins). Traits and terms with identical sets of confidently-linked genes were merged for each source of evidence. Genes less confidently associated with each of the merged traits (OTAR score < 0.5) were collected in similar fashion. Evidence was analogously collected from OTAR and processed for IMPC mouse phenotypes (http://www.mousephenotype.org/) (Dickinson et al. 2016). Gene Ontology terms were filtered for not being obsolete and for having a description, and then annotated with associated genes using the QuickGO annotations from EBI (date = aug. 2024). Additionally, the GO terms were annotated with associated genes reported by UniprotKB (date = nov. 2024) through their Rest API. Both sets of associated genes were merged and gene identifiers were converted to Ensembl identifiers (Ensembl Biomart, August 27, 2024). Finally, the GO terms with identical gene sets were merged for each GO namespace.

### Quantifying discordance of GWAS and ClinVar (Fig. 1AB)

Variants from GWAS and ClinVar were grouped using the variantFunctionalConsequenceId and the Sequence Ontology tree (Eilbeck et al. 2005), aggregating the ontology tree into exon, intron, splicing, intergenic, and other structural variants. Similarity of evidence between GWAS and ClinVar was quantified using gene-disease associations that included indirect evidence (’associationByDatasourceIndirect’). The percentages of genes linked through each of the sources of evidence was computed for traits that had evidence from both GWAS and ClinVar (1,412 traits). Finally, traits were filtered for having paired EFO and ICD10 codes in FinnGen (Kurki et al. 2023) and annotated with their prevalence (287 traits).

### Quantifying discordance across sources of evidence (Fig. 1CD)

Similarities between selected sources of evidence were quantified as follows. Similarity of traits was scored as the number of traits shared between two sources of evidence, expressed as a percentage of the possible maximum. Similarity of linked genes was scored as the maximum possible percentage of shared genes, averaged over the traits that had shared linked genes. Gene-level and protein-level characteristics were defined as the average log2 protein abundance (PaxDB integrated whole-organism) (Huang et al. 2023), the average gene essentiality (DepMap CRISPRGeneEffect; https://depmap.org/portal) (DepMap 2024), the log2-transformed median-confidence interactions (STRING score >= 0.4) (Szklarczyk et al. 2019), and the median GC content (Ensembl Biomart). Gene symbols and other identifiers were converted to Ensembl IDs (Ensembl Biomart). These characteristics were then compared between the confidently-linked genes for a source of evidence (OTAR evidence score >= 0.5) and all other genes for which the property was quantified (two-sided Mann-Whitney U test - p-values were FDR corrected through the Benjamini Hochberg procedure).

### Computing network embedding of proteins

Evidence for protein-protein interactions were collected from the Open Targets Genetics Portal (version 24.09). The evidence was filtered for medium-confidence physical interactions from IntAct (score >= 0.45; following (Villaveces et al. 2015)), medium-confidence interactions in STRING (Szklarczyk et al. 2019) (score >= 0.4), and interactions from Signor (Lo Surdo et al. 2023) or Reactome (Milacic et al. 2024). Interactions were further filtered for proteins having an Ensembl identifier (’ENSG’ IDs) and (for network analysis purposes) for being part of the main connected component of the network. The final network contained 19,432 genes with 1,000,657 interactions. Rewired networks were generated by sequentially swapping the interactors of random pairs of edges, thereby preserving each node’s degree. The number of swaps equalled 10x the number of protein interactions for each rewired network. The resulting 100 rewired networks on average shared 38,975 interactions with the original protein interaction network, similar to the number of shared interactions found for interaction networks generated through the configuration model following the same degree distribution (37,682 interactions, n=10).

Network embeddings of genes were then created following the node2vec algorithm (Grover and Leskovec 2016) (`gensim` package) for each of the rewired networks and the protein interaction network. Specifically, 30 random walks of length 100 were generated starting from each node in the network (p=1, q=1). These random walks were then used to train a n=128 dimensional skip-gram model (window_size = 10; 5 epochs).

### Computing network scores for traits and terms

The node2vec embeddings of proteins for the protein interaction network and rewired networks were used to generate network scores for traits and terms. Specifically, the skip-gram model for each network (from ‘Computing network embedding of proteins’) was first used to create a transition matrix that contained the probabilities to observe each protein during a random walk involving any other protein (i.e., using the model’s hidden state and each protein’s embedding to obtain a score vector for other proteins that is normalized with the sum of scores to obtain probabilities (Grover and Leskovec 2016)). Probabilities for genes to be involved with disease through network *k* were then scored using the probability to see at least one disease gene (i.e., 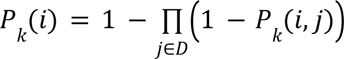 for the set of disease genes *D*). Network scores were then obtained by log-normalizing the probabilities for the protein interaction network with the geometric mean of the probabilities from the rewired networks (i.e., the difference of the log-transformed probabilities for the protein interaction network and the average of the log probabilities for the rewired networks). The trained models for computing network scores are available through BioStudies (S-BSST2641).

### Sub-samples of traits by sampling confidently-linked genes

To generate samples, traits and terms were filtered for having at least n=20 confidently-linked genes (n=5,857 traits and terms). For each of these traits and terms, n=100 samples were created by randomly sampling 75% of the confidently-linked genes. Specifically, all confidently-linked genes were randomly ordered in a list, of which three copies were concatenated and then split into four equally-sized parts. This procedure created four unique samples that each contained 75% of the confidently-linked genes and together contained each confidently-linked gene three times - ensuring that the confidently-linked genes were sampled equally frequently. Leftover genes were ignored (i.e., when the number of confidently-linked genes was not a multiple of four). Network scores were then computed for each of these samples of all of the traits and terms (following ‘Computing network scores for traits and terms’).

### Computing latent representations for traits and terms

A standard Variational Autoencoder (VAE) was used to create latent representations of the network scores for traits and terms. The encoder consisted of a fully connected hidden layer with leaky ReLU activation, followed by fully connected layers for the mean and variance and finally the sampling. The decoder consisted of a fully connected hidden layer with leaky ReLU, followed by a final fully connected layer. During training, the loss consisted of equally weighted reconstruction loss (measured by the mean squared error) and Kullback–Leibler divergence. The VAE was trained using the network scores of 38,826 traits and terms (from ClinVar, GWAS, Orphanet, GO terms and mouse phenotypes). For each trait and term with at least 20 confidently-linked genes, its first four sub-samples (from ‘Sub-samples of traits by sampling confidently-linked genes’) were collected as an independent test set for the VAE (n=23,428 samples). During training, the performance was evaluated for every epoch on both the training and test data, evaluating the model’s KL loss, reconstruction loss, and the person and spearman correlations for samples before encoding and after decoding. The model was trained using the Adam optimizer, a batch size of 500 and a learning rate of 0.000033. The dimensionality of the latent space was chosen to have linearly independent latent dimensions (quantified through PCA) while maximizing the recovery of the test samples (quantified through an independent metric not used for training - the Spearman correlation between network scores and their reconstruction) (also see Supplementary Figs. 1-2). The final model was trained for 15 epochs using n=128 latent dimensions and hidden layers of size 1,024. For the final model and the independent test samples, these correlations were on average 0.926 (Pearson) and 0.944 (Spearman). Finally, the network scores of traits and terms from all sources of evidence and the sub-samples of ClinVar traits were encoded with the trained VAE using the output of the ‘mean’ layer from the encoder. The trained VAE is available through BioStudies (S-BSST2641).

### Categorizing traits and terms

Relational information was obtained for human traits and diseases (from Open Targets; version 24.09), GO terms (from go-basic; date = 24-06-17), and mouse phenotypes (mouse genome informatics (MGI) consortium; 25-01-30). Parental lineages were collected for each of these traits and terms using the respective ontology trees. Siblings were determined through traits and terms sharing a parent. Lineages were merged for traits and terms that had identical sets of linked genes in the interaction network (see ‘Evidence for associations of genes with traits and terms**’**). The resulting table described the direct parents, children and siblings of each trait and term according to the ontology trees, merged for identical sets of linked genes. Analogously, these lineages yielded more distant parental and child traits and terms further upstream and downstream in the ontology trees. Traits and terms were then assigned broad categories using the top-level annotations from the ontology lookup service and manual curation (EMBL-EBI; e.g., http://www.ebi.ac.uk/efo/EFO_0000408). Here, categories were filtered for having at least 10 children with network scores. Large categories (e.g., identifiers having >1,000 children) were sub-divided into smaller ones through lower-level annotations, and umbrella terms were additionally split into respective categories (e.g., ‘musculoskeletal’ into ‘musculature’ and ‘skeletal’). These categories were then harmonized across ontology tables through manual curation, assigning 306 top-level identifiers from 6 different ontologies (MONDO, EFO, Orphanet, HP, MP and GO) into 34 broad categories of traits and terms (Supplementary Table 1). Finally, the table of lineages was used to annotate the network scores for traits and terms using the lineages and annotations of broad categories. Annotations of traits and terms were not unique (e.g., a trait could be assigned all of ‘cardiovascular’, ‘heart’, ‘vascular’, ‘hematopoietic’, and ‘musculature’). Among the traits and terms with network scores, this methodology assigned broad categories to 92% of ClinVar traits, 75% of GWAS traits, 96% of mouse phenotypes and 58% of GO terms (Supplementary Table 2).

### Comparing distances of similar traits with unrelated traits (Fig. 2B-C)

Euclidean distances were computed between the latent representations of traits (Burden, GWAS, ClinVar and Orphanet) and other traits or terms (GO terms, mouse phenotypes, drugs). Here, GO terms were selected for having at least 5 linked genes (6,913 terms). These distances were then used for further analyses. Here, traits from Burden, GWAS and ClinVar were filtered for having at least 5 confidently-linked genes (see ‘Evidence for associations of genes with traits and terms**’**). Jaccard indices were computed between all pairs of these traits across all three sources of evidence, and compared with the distances. For a given trait, all other traits were then labeled as ‘same’, ‘similar’ or ‘other’, within and between sources of evidence. Specifically, for a given trait, a trait was labeled as ‘same’ if at least one of their identifiers was identical (see ‘Evidence for associations of genes with traits and terms’). Similarly, a trait was labeled as ‘similar’ if they were siblings according to the ontology tree, while a trait was labeled as ‘other’ if it was not the same, a parent, sibling or child of the given trait in the ontology tree (see ‘Categorizing traits and terms’). Finally, for a given trait and sources of evidence, the average distance was computed to traits that were labeled as same, similar or other. Traits without siblings were ignored when comparing between these categories.

### Mapping the landscape of traits and terms (Fig. 2D-E)

Latent representations of all 38,826 traits and terms were used to fit a 2-dimensional UMAP. The annotation of terms and traits was simplified for visualization purposes as follows. Traits and terms were first annotated with n=18 of the broad categories (from ‘Categorizing traits and terms’), taking for each trait and term the most likely category as estimated from probabilities of gaussian kernel densities fitted to the UMAP coordinates of terms and traits assigned to each category (annotations for 23,595 traits and terms). Outliers in the UMAP were not visualized by only including traits and terms whose UMAP coordinates were at most one unit outside of the range of coordinates for traits and terms that had an annotation - this excluded n=951 of traits and terms (<2.5%), none of which had an annotation. Finally, traits and terms were only colored according to their most likely annotation if its estimated probability exceeded the median probability from all categories (63.8% of all visualized traits and terms - n=15,043). The same UMAP coordinates were used to color all traits and terms according to their source of evidence.

### Comparing distances of drugs to disease (Fig. 2G)

Molecules were linked to diseases through molecule-disease indications with maximum clinical phase from ChEBML (https://www.ebi.ac.uk/chembl/; via Open Targets). Protein-molecule associations were collected from the STITCH database (molecules acting on human proteins; https://stitch-db.org/) and filtered for confident associations (STITCH score >= 0.7). Targets were filtered for having a network embedding (’Network embedding of proteins’), after which molecules with identical target protein sets were merged. Network scores and latent representations of molecules were created (following ‘Computing network scores for traits and terms’ and ‘Computing latent representations for traits and terms’). These network scores and latent representations of molecules were then used for further analysis. Specifically, drugs were filtered for having at least n=5 target proteins (1,201 drugs; following ‘Comparing distances of similar traits with unrelated traits’) and for having reached clinical phase >= 4 for at least one disease (n=454 drugs; ‘approved drugs’). Similarly, diseases were filtered for having at least one approved drug, and distances were computed between all remaining drugs and traits for each source of evidence. Finally, the average distances of drugs to their approved diseases and to all other diseases with approved treatment were computed for each source of evidence.

### Consensus clustering of the interaction network into protein modules

The protein interaction network was clustered into functional protein modules using an ensemble Leiden community-detection procedure with edge subsampling from the network and recursive subdivision of large clusters. Both the edge subsampling and varying the resolution for clustering were applied to increase the stability of the clustering of proteins, while recursive subdivision of clusters allowed for the recovery of finer substructures in denser regions of the interaction network. First, 1,000 subgraphs were generated by sampling a fraction of edges from the interaction network. Here, the fractions of sampled edges were drawn from the Beta(16, 4) distribution (with mean 0.8). Leiden clustering was then performed on these subgraphs, using resolution parameters sampled from the Beta(2, 14) distribution (with mean 0.125) (`python-igraph` package, version 0.11.8). Finally, for each clustered subgraph, the clusters exceeding a threshold of 200 nodes were recursively re-clustered following the same procedure with their respective subgraphs, till they were below the threshold or up to a maximum recursion depth of 10 steps. The clustered subgraphs were then used to construct a pairwise agreement matrix describing the fraction of subgraphs for which each pair of nodes was in the same cluster. A consensus graph was then defined by connecting all pairs of nodes whose agreement fraction was larger than 0.3, using the agreement fractions as edge weights. This weighted consensus graph was then clustered through Leiden clustering (resolution 0.045 - chosen as a trade-off between average module size and number of genes in very small (n<4) modules) (Supplementary Table 4).

### Associating protein modules to terms and traits

The protein interaction network was clustered into functional modules following ‘Consensus clustering of the interaction network into protein modules’. Protein modules were filtered for containing at least 5 proteins from the interaction network. For each of these modules, the number of overlapping genes was computed with all traits and terms from each of the sources of evidence (ClinVar, Burden, GWAS, GO terms and mouse phenotypes). The association of modules with terms and traits was tested through gene-set enrichment analysis (GSEA - R `fgsea` package version 1.31.2; eps = 0) using the network scores (from ‘Computing network scores for traits and terms’). Finally, Modules were associated to traits and terms by requiring at least one shared gene and an enrichment of the module’s genes among the top ranked proteins according to the network scores (Benjamini-Hochberg (BH)-adjusted p-values < 0.001; GSEA). This approach yielded 210,392 confident associations between 889 protein modules and 34,692 traits and terms (0.68% of the possible associations) (Supplementary Table 5).

### Comparing convergence of similar traits with unrelated traits (Fig. 3B-C)

Convergence between pairs of traits and terms was quantified through the Jaccard index of their confidently associated protein modules (from ‘Associating protein modules to terms and traits’), both within and across sources of evidence (for ClinVar, Burden, GWAS). Here, the traits and terms were filtered for having at least 5 confidently-linked genes. Convergence between similar and unrelated traits was then quantified analogous to ‘Comparing distances of similar traits with unrelated traits’.

### Logistic model for scoring the likelihood for traits to be related (Fig. 3D)

The Euclidean distances between the latent representations of ClinVar traits were used to compute the average distance of a trait to all similar traits (siblings in the ontology tree) or with other traits (not the same trait according to overlapping identifiers, not a parent, sibling or child of the given trait in the ontology tree - see ‘Categorizing traits and terms’). Traits were then filtered for having at least one similar trait, and a logistic model was fitted to the distances using the relations ‘similar’ or ‘other’ as a label (`LogisticRegression` from `scikit-learn`; using balanced class weights and an intercept - model accuracy = 0.59). This model yielded the cut-off latent distance for which ClinVar traits have probability 0.5 to be related (distance = 24.98).

### Quantifying enrichment of clusters with broad categories of traits (Fig. 3D)

The ClinVar traits with network scores were filtered for the n=570 traits that also had network scores from GWAS but were not neoplasms (‘Categorizing traits and terms’). These traits were then clustered through complete-linkage clustering with the Euclidean distance of their latent representations from the ClinVar network scores. Clusters were filtered for containing at least 8 traits and having a maximum distance between any pair of their traits below the distance cut-off (n=125 clusters; from ‘Logistic model for quantifying the likelihood for traits to be related’). Enrichment for these clusters with traits from each of the broad categories was quantified through a one-sided Fisher exact test, using the other clustered traits as background. Clusters were required to be significantly enriched with at least one broad category (BH-adjusted p-value < 0.05). Overlaps between enriched clusters were resolved through a greedy approach by selecting the cluster with the strongest enrichment (highest odds-ratio) for one of the broad categories and discarding all overlapping clusters until no clusters with enrichment were left. Finally, the selected clusters with strongest enrichment were substituted with “parent clusters” (any enriched cluster containing the selected cluster) if the parent fully contained at least two of the selected clusters with enrichment for the same broad category and if the parent did not contain any selected cluster enriched for other broad categories of traits. These parent clusters were selected by taking the parent with the most selected clusters, the strongest enrichment for the broad category, and for being the smallest parent that is an enriched cluster itself. This procedure grouped n=263 (46.1%) of the ClinVar traits with GWAS evidence into n=22 clusters from n=14 broad categories.

### Quantifying enrichment of modules linking to trait clusters (Fig. 3E)

Enrichment of trait clusters (from ‘Quantifying enrichment of clusters with broad categories of traits’) with protein modules (from ‘Consensus clustering of the interaction network into protein modules’) was computed through a one-sided Fisher exact test with the cluster’s ClinVar traits associated to a given protein module (see ‘Associating protein modules to terms and traits’). The ClinVar traits with GWAS evidence but not in that cluster and with a different annotation than the cluster’s broad category were used as the background (there were several trait clusters enriched with the same broad category). Trait clusters were linked to a protein module if at least half the traits in the cluster were associated to that module and the cluster was enriched with associations to the module (BH-adjusted p-value < 0.01). This approach identified n=301 links of n=135 protein modules to the n=22 trait clusters.

### Quantifying convergence between ClinVar and GWAS traits linking to modules (**Fig 3F**)

Module-trait associations (from ‘Associating protein modules to terms and traits’) for traits with network scores from ClinVar and GWAS were aggregated at the level of the trait clusters (from ‘Quantifying enrichment of clusters with broad categories of traits’) by quantifying whether at least one trait in the cluster linked to a given module for each source of evidence. Convergence of the sources of evidence for linking to the same modules was then computed as the odds-ratio of a one-sided Fisher exact test.

### Labeling modules through enrichment with genes from GO terms

Protein modules were tested for enrichment with genes from GO terms (from ‘Evidence for associations of genes with traits and terms’) through one-sided Fisher exact tests using all genes linked to any GO term and all genes linked to any module as backgrounds. The modules were then annotated with representative names from the top 5 GO terms having the strongest enrichment for the respective modules (requiring at least odds-ratio >= 10 and BH-adjusted log p-values > 10) (Supplementary Table 7).

### Visualizing interaction network for sensorineural hearing loss (Fig. 3G-H)

The n=18 modules linked to the trait cluster of n=8 auditory traits were labeled through the GO terms (from ‘Labeling modules through enrichment with genes from GO terms’). The stereocilium module was then selected as both the only module linked to all auditory traits through at least one source of evidence and one of the modules linked to the most traits through both sources of evidence. Sensorineural hearing loss (SNHL) was selected in the same manner. All proteins from the stereocilium module were annotated for being weakly- or confidently-linked SNHL genes from ClinVar and GWAS, being linked to SNHL phenotype through mouse knockouts, and for having supporting evidence from at least 5 of the other GWAS and ClinVar traits in the trait cluster. Specificity of prioritization by sources of evidence was performed through comparison of the network scores of SNHL for curated gene sets with paired t-tests. The visualized network was constructed as follows. The protein interaction network was filtered for proteins in the stereocilium module, and for interactions having moderate-confident scores (from ‘Computing network embedding of proteins’). Self-interactions were added for all proteins in the module (score = 1), and the interactions were sorted by score, removing duplicates with lower scores. The interaction network of the module was then simplified for visualization by sequentially removing all interactions (with scores < 0.95) that were not necessary to keep the network fully connected, in ascending order of interaction scores. Finally, the size of the nodes was defined as the average of the network scores for SNHL from GWAS and ClinVar, while the color of the nodes was defined as the difference in network scores between GWAS and ClinVar relative to the average (‘network bias’). This network bias was clipped to 50% (one protein that was not a confidently-linked SNHL gene had a more extreme value (SPTAN1)). Gene identifiers were converted to HGNC symbols through Ensembl Biomart (see ‘Evidence for associations of genes with traits and terms’) (Supplementary Table 8).

### Quantifying recovery of disease genes with network scores (Fig. 4A)

Recovery of linked genes was quantified through the Area Under the Receiver Operator Characteristic (AUROC) curve and the average precision-recall (PR) using the network scores for the sub-samples of traits (from ‘Sub-samples of traits by sampling confidently-linked genes’ - also see Supplementary Fig. 10). First, we removed the 75% of confidently-linked genes that were used for computing the network scores. Recovery of linked genes was then scored using the other 25% of withheld confidently-linked genes or all of the weakly-linked genes (see ‘Evidence for associations of genes with traits and terms’). Specifically, positives were defined as the weakly- or confidently-linked genes for a given trait, using all other genes with a network score as negatives. The AUROC scores were z-scored using sampled sets of disease genes that followed the same frequency distribution for the source of evidence of the respective trait. Specifically, the frequency distribution for linked genes was defined as the number of times each gene was (weakly- or confidently-)linked to diseases for a given source of evidence. Disease genes were then randomly drawn for each trait using the respective frequency distribution, generating n=100 samples of disease genes for each trait and source of evidence. Genes (weakly- or confidently-)linked to the traits were removed from the respective frequency distribution before sampling, and the number of sampled genes matched the linked genes for that trait at the given confidence level if sufficient genes were available in the distribution. Traits were omitted if the samples of disease genes had a median Jaccard index that exceeded 0.1 (i.e., the frequency distribution had too few genes or was too skewed to yield substantially different samples of disease genes for z-scoring the AUROC; e.g., for EFO_0001444 (measurement) from Burden and GWAS). The resulting samples of disease genes tested the specificity of network scores to recover trait-linked genes compared to other disease-linked genes for the same source of evidence by matching their frequencies and sample size. Finally, these sampled disease genes were used to score the AUROC for each of the traits using the network scores of the trait’s sub-samples. AUCs and average PRs were computed using the `roc_auc_score` and `average_precision_score` functions from the `scikit-learn` package respectively.

### Classifying traits as nearest neighbors

Distances between traits and terms were computed as the Euclidean distances between the latent representations of the network scores from traits and terms. A logistic model was then fitted to the distances between similar and unrelated traits across sources of evidence (ClinVar, Burden and GWAS - analogous to ‘Logistic model for scoring the likelihood for traits to be related’). Here, for each trait, the average distances were computed to similar traits (siblings in the ontology tree; from any source) and to other traits (see ‘Categorizing traits and terms’), which were used as positives and negatives respectively (model accuracy = 0.67). We then took the average distance between traits and other traits from any source of evidence as the cut-off distance for traits to no longer be considered close neighbors (distance = 15.68; probability = 0.41 to be similar according to the logistic model). Close neighbors were then defined as the traits or terms from a given source of evidence whose latent representations were within this cut-off distance from a trait of interest.

### Quantifying recovery of disease genes by neighboring traits (Fig. 4C)

The top 5 nearest traits or terms from each source of evidence (from ‘Classifying traits as close neighbors’) were selected for all of the n=100 sub-samples of n=704 ClinVar traits (from ‘Sub-samples of traits by sampling confidently-linked genes’) using the Euclidean distances between the latent representations. These nearest traits or terms were defined as the ‘close neighbors’ and could not be the same trait as the respective sub-samples. The latent representations for these close neighbors were averaged and then decoded using the VAE (from ‘Computing latent representations for traits and terms’) if 5 such neighbors were available. The reconstructed network scores from the averaged close neighbors were then used as predicted scores for the sub-samples of ClinVar traits from a given source of evidence. Recovery of trait-linked genes was then quantified by first removing the 75% of trait-linked genes used for computing the latent representation of a sub-sample and then recovering the 25% remaining trait-linked genes (following ‘Quantifying recovery of disease genes with network scores’, also see ‘Sub-samples of traits by sampling confidently-linked genes’ and Supplementary Fig. 10). Traits from Burden were omitted as neighbors because their low number and larger distances to the ClinVar traits would be restrictive for the traits having sufficiently close neighbors.

### Recovery of disease genes by more distant traits (Fig. 4C)

Distant neighborhoods for sub-samples of ClinVar traits were generated by first sampling distant traits (or terms) and then selecting the nearest neighbors for these distant traits. Specifically, distant traits were sampled for each of the n=100 sub-samples for n=704 ClinVar traits (from ‘Sub-samples of traits by sampling confidently-linked genes’) by requiring the Euclidean distances between their latent representations to at least 25% above the distance cut-off for traits to no longer be close neighbors (distance = 19.6; probability = 0.19 to be similar traits - from ‘Classifying traits as close neighbors’). This generated n=100 distant traits or terms from each source of evidence for each of the sub-samples of ClinVar traits. Next, for each of these distant traits (or terms), the top 4 nearest neighbors were selected from the same source of evidence that were still at least the cut-off distance (15.68) away from the respective sub-sample. Together, the distant trait and its 4 nearest neighbors created a distant neighborhood of traits for the sub-sample that were all at least the cut-off distance removed from the sub-sample, thereby creating n=100 sampled distant neighborhoods for each of the sub-samples of the ClinVar traits. Predicted scores for the distant neighborhoods and recovery of the trait-linked genes were then scored as before (analogous to ‘Quantifying recovery of disease genes by neighboring traits’).

### Prioritizing candidate genes for rare diseases through network analysis (Fig 4D-E)

The top n=5 nearest latent neighbors from each source of evidence within the cut-off distance (from ‘Classifying traits as nearest neighbors’) were selected for familial Long QT Syndrome from ClinVar (fLQTS - MONDO_0019171), including drugs (from ‘Comparing distances of drugs to disease’). Neighbors from all sources of evidence were clustered using their latent representations and annotated with overlaps in disease-linked genes and their distances to fLQTS (from ClinVar). Next, the neighbors from each source of evidence were averaged by their latent representation and decoded to obtain reconstructed network scores for each source of evidence. Genes were selected for ranking among the top 100 in the predicted network scores by the nearest neighbors from at least two sources of evidence. The protein interaction network was then filtered for highly confident interactions (OTAR interaction score >= 0.95) between these filtered genes and including the confident fLQTS-linked genes. The predicted scores from the nearest neighbors were averaged over the sources of evidence and sorted by rank. Ranks were capped at 1,000 and log-transformed to obtain an average rank score from the nearest neighbors used for the size of the nodes (one protein had a more extreme value (CALM2)). Following the same analysis, the network scores of fLQTS from ClinVar were used to obtain a rank score for coloring the nodes. Gene identifiers were converted to HGNC symbols through Ensembl Biomart (see ‘Evidence for associations of genes with traits and terms’). Consensus RNA expression data from The Protein Atlas (https://www.proteinatlas.org/) was used to compute the heart-specificity of genes in the heart muscle (requiring z-score >= 2 after z-score across tissues and requiring gene expression to be among the top 5% of expressed genes in the tissue) (Supplementary Table 9).

### Systematically prioritizing candidate genes for rare diseases

Genes were systematically prioritized for rare diseases as before (from ‘Prioritizing candidate genes for rare diseases through network analysis’). To do so, ClinVar traits were filtered for having an annotation (from ‘Categorizing traits and terms’), not being neoplasms, and having at least n=3 confidently-linked genes. Nearest neighbors from ClinVar, GWAS, GO terms and Mouse phenotypes were collected for the remaining n=2,141 ClinVar traits. Specifically, n=5 neighbors were required to be within the cut-off distance (from ‘Classifying traits as nearest neighbors’) for each source of evidence and at least two of these neighbors (for ClinVar or GWAS) were required to share an annotation with the respective trait. ClinVar traits were further filtered for having an AUROC >= 0.7 when recovering their confidently-linked genes with the predicted scores of the nearest neighbors from each source of evidence (Supplementary Table 10). Targets were then prioritized for the n=477 remaining ClinVar traits. For each of these traits and as before (from ‘Prioritizing candidate genes for rare diseases through network analysis’), proteins were filtered for ranking among the top n=100 proteins according to the predicted scores from at least two sources of evidence. For further filtering, genes were then required to have weaker prior evidence (OTAR scores between 0.25 and 0.5 for the respective trait) (Supplementary Tables 11-12).

## DATA AVAILABILITY

All processed, analyzed and generated data from this study are publicly available (BioStudies S-BSST2641). Evidence for gene-trait associations was collected from the Open Targets (OTAR) genetics portal (version 24.09; https://platform.opentargets.org/) (Ghoussaini et al. 2021; Mountjoy et al. 2021). The protein-protein interaction network was collected from Open Targets and includes evidence from STRING (Szklarczyk et al. 2019), IntAct (Del Toro et al. 2022), Reactome (Milacic et al. 2024) and Signor (Lo Surdo et al. 2023). Evidence for mouse phenotypes was collected from IMPC through OTAR (http://www.mousephenotype.org/) (Dickinson et al. 2016). Evidence for molecule-disease indications were collected from ChEMBL (https://www.ebi.ac.uk/chembl/; via Open Targets), and protein-molecule associations were collected from the STITCH database (https://stitch-db.org/). Gene Ontology terms and linked genes were collected using the Uniprot rest API (https://www.uniprot.org) and EBI (https://www.ebi.ac.uk/QuickGO). Gene and protein identifiers and symbols were converted with Ensembl biomart (https://www.ensembl.org). Consensus RNA expression data was collected from the Protein Atlas (https://www.proteinatlas.org), protein abundance data was obtained from PaxDB (integrated whole-organism) (Huang et al. 2023), and gene essentiality was collected from DepMap (CRISPRGeneEffect; https://depmap.org/portal) (DepMap 2024).

## CODE AVAILABILITY

All code for data processing, analysis and the figures in this study is publicly available as Jupyter notebooks (BioStudies S-BSST2641).

## SUPPLEMENTARY FIGURES

**Figure S1.**
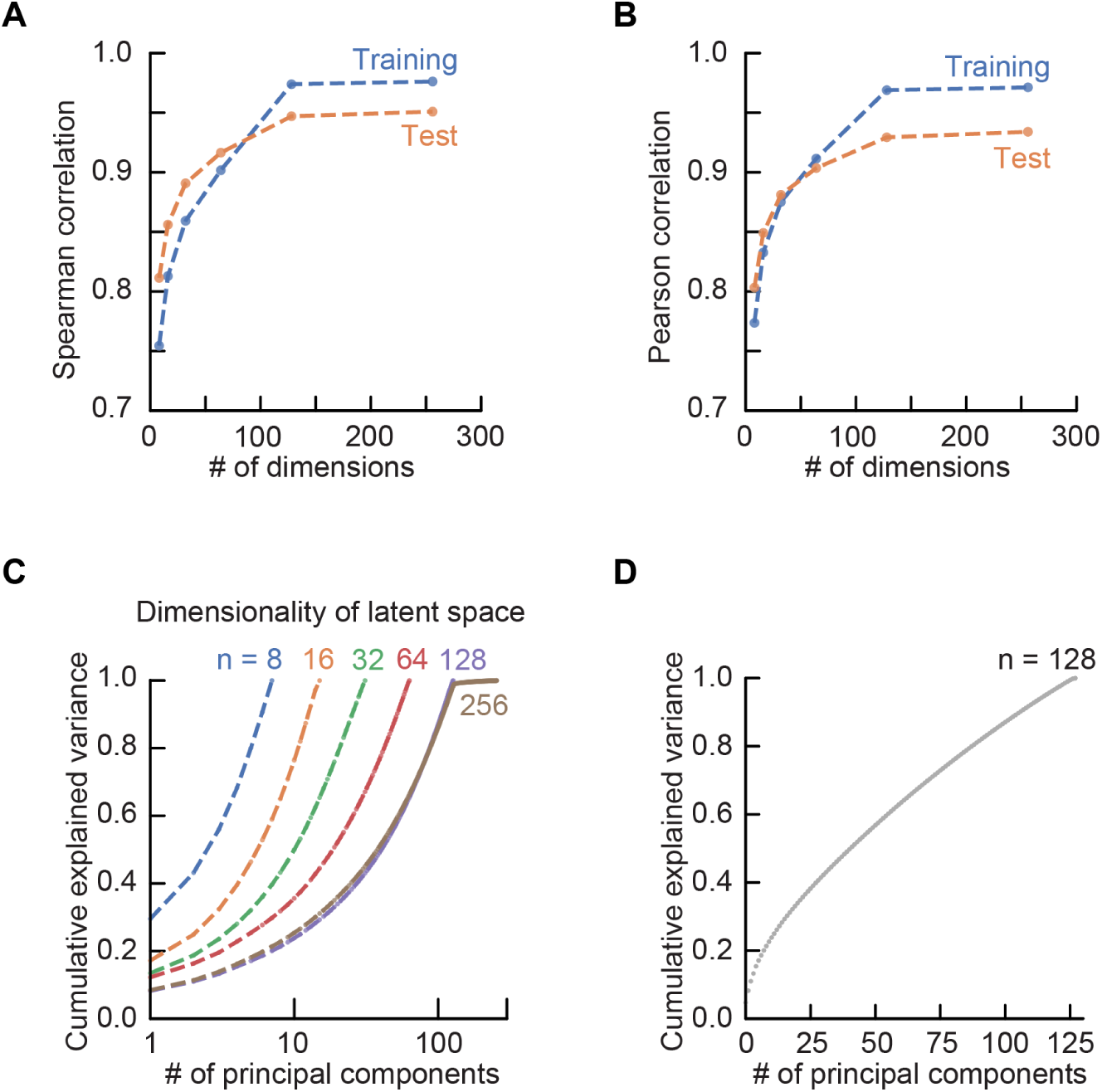
Latent dimensionality dictates recovery of network scores from latent space (Related to. Fig. 2A). VAE’s were trained with varying latent dimensionality using the network scores of n=38,826 traits and terms (ClinVar, GWAS, Burden, Orphanet, GO terms and mouse phenotypes - Methods). Models were trained for n=30 epochs. **(A-B)** Shown are the average Spearman **(A)** and Pearson **(B)** correlation coefficients of the input and reconstructed network scores as a function of the dimension of the VAE’s latent space (n = 8, 16, 32, 64, 128 (all having hidden layers of size 1,024) and n = 256 (hidden layers of size 2,048)). Performance of these models was quantified for both the training samples (blue) and n=23,428 independent test samples obtained from computing network scores on samples of confidently-linked genes (orange; Methods). Both the Spearman and Pearson correlation coefficients are independent and appropriate metrics for evaluating the model’s performance as these metrics are not used for the training loss of the VAE and because both metrics are important for recovering trait-linked genes by their rank or score in the network scores. Both Spearman and Pearson correlation coefficients increase with an increasing number of latent dimensions, plateauing at n>=128 dimensions (e.g., ρ = 0.947 for n=128 and ρ = 0.951 for n=256; Spearman correlation on the test set). Next, the latent representations of all traits and terms was used to determine the explained variance by the latent dimensions through a Principal Component (PC) analysis. **(C-D)** Shown is the cumulative explained variance as a function of the number of PCs for the various dimensionalities of the latent space **(C)** and in particular for n=128 dimensions **(D)**. This analysis demonstrates that the cumulative explained variance for the first 128 principal components in a 256-dimensional latent space was 0.984, with the last 128 principal components explaining a negligible variance in the latent representations of traits and terms. Together, these observations demonstrate that reconstruction of network scores - as quantified through correlation - improves up to n=128 latent dimensions, with larger latent spaces not substantially improving performance and not explaining more variance in the samples compared to smaller latent spaces. Overall, we used a n = 128 dimensional latent space for the VAE to compress the network scores of traits and terms.

**Figure S2.**
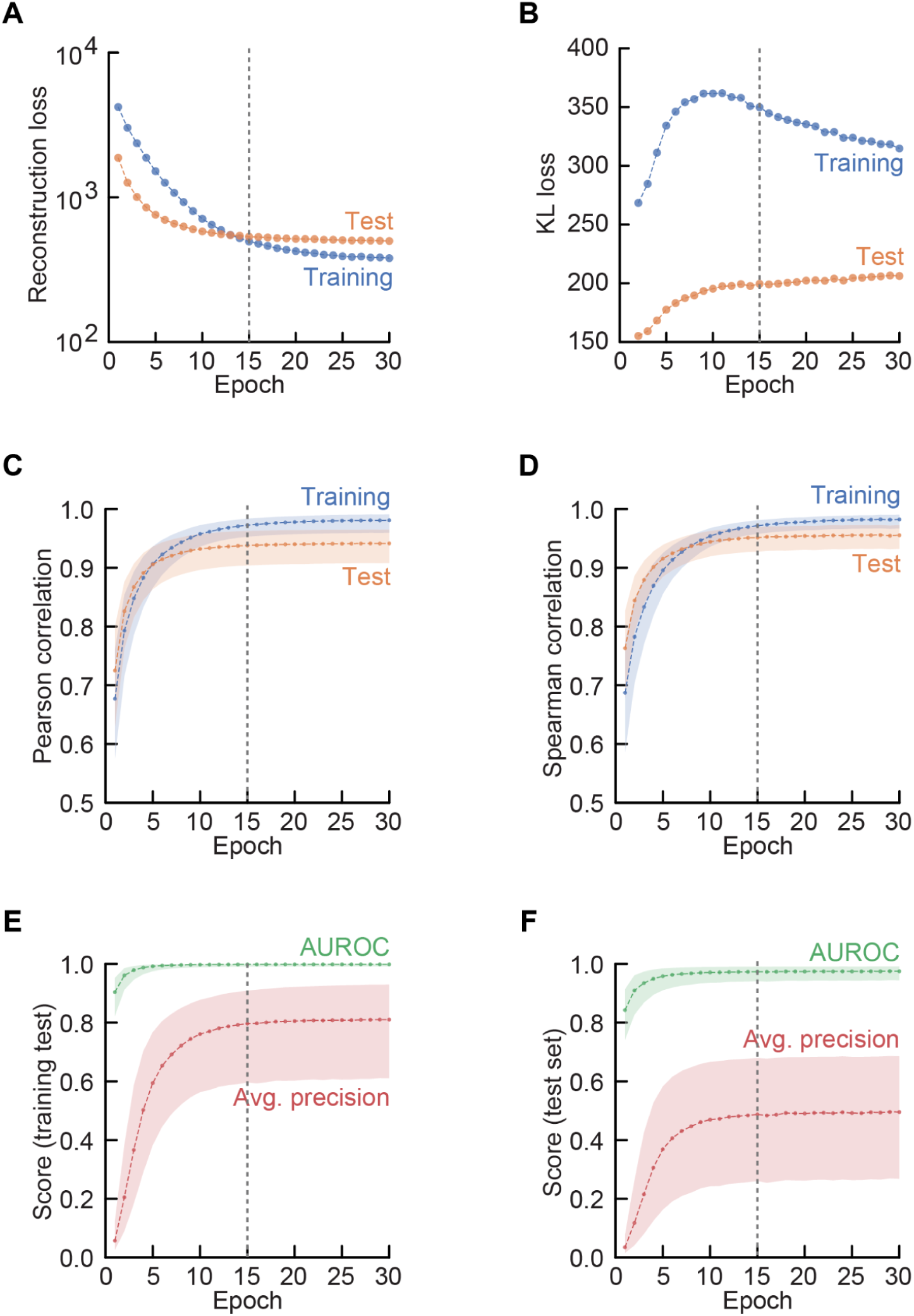
Training a VAE for latent representations of traits and terms (Related to Fig. 2A). A VAE was trained with n=128 latent dimensions following Supplementary Fig. 1 (also see Methods). The model was trained for n=30 epochs. **(A-B)** Shown is the Reconstruction loss as measured by the mean squared error with the network scores **(A),** and the KL loss quantifying the Kullback–Leibler divergence **(B),** for both the training samples (blue) and the test samples (orange; Supplementary Fig. 1 and Methods). Both the reconstruction loss and the KL loss did not substantially improve after 15 epochs, while the training losses continued to decrease. **(C-D)** To test the performance of the model as a function of the number of epochs, we quantified the correlation coefficients between the network scores and their reconstruction. Shown are the Spearman **(C)** and Pearson **(D)** correlation coefficients for the training (blue) and test (orange) samples as a function of the number of epochs. Dots show the median, shaded area shows first and third quantiles of the samples. We found that the correlation coefficients for the reconstructed network scores had plateaued after n=15 epochs (grey dotted line), with a median Spearman correlation of ρ = 0.951 (0.93 for Q1) compared to ρ = 0.955 after n=30 epochs. **(E-F)** Finally, we quantified the ability of the reconstructed network scores for recovering the top ranked genes. Specifically, we determined whether the reconstructed scores could recover the top n=100 ranked genes of the network scores for each trait and term, as quantified by the Area Under the Receiver Operating Characteristic curve (AUROC) and Average Precision score (AP). Shown are the AUROC (green) and AP scores (red) for the training samples **(E)** and the test samples **(F)** as a function of the number of epochs. Dots show the median score, shaded area shows the first and third quantiles of the samples. As expected, the recovery of top scoring genes for training samples outperforms the recovery of top scoring genes for test samples and plateaus at n=15 epochs (grey dotted line). Specifically, after n=15 epochs, the reconstructed scores from the VAE recover the top scoring genes with a median AUROC of 0.974 (0.942 for Q1) and a median AP of 0.487 (0.260 for Q1), compared to a median AUROC of 0.975 and a median AP of 0.495 after n=30 epochs. These analyses demonstrate that the performance of the reconstructed scores from the trained model does not substantially improve after n=15 epochs. We therefore chose to train the VAE used for creating latent representations of traits and terms for n=15 epochs (Figs. 2-4).

**Figure S3.**
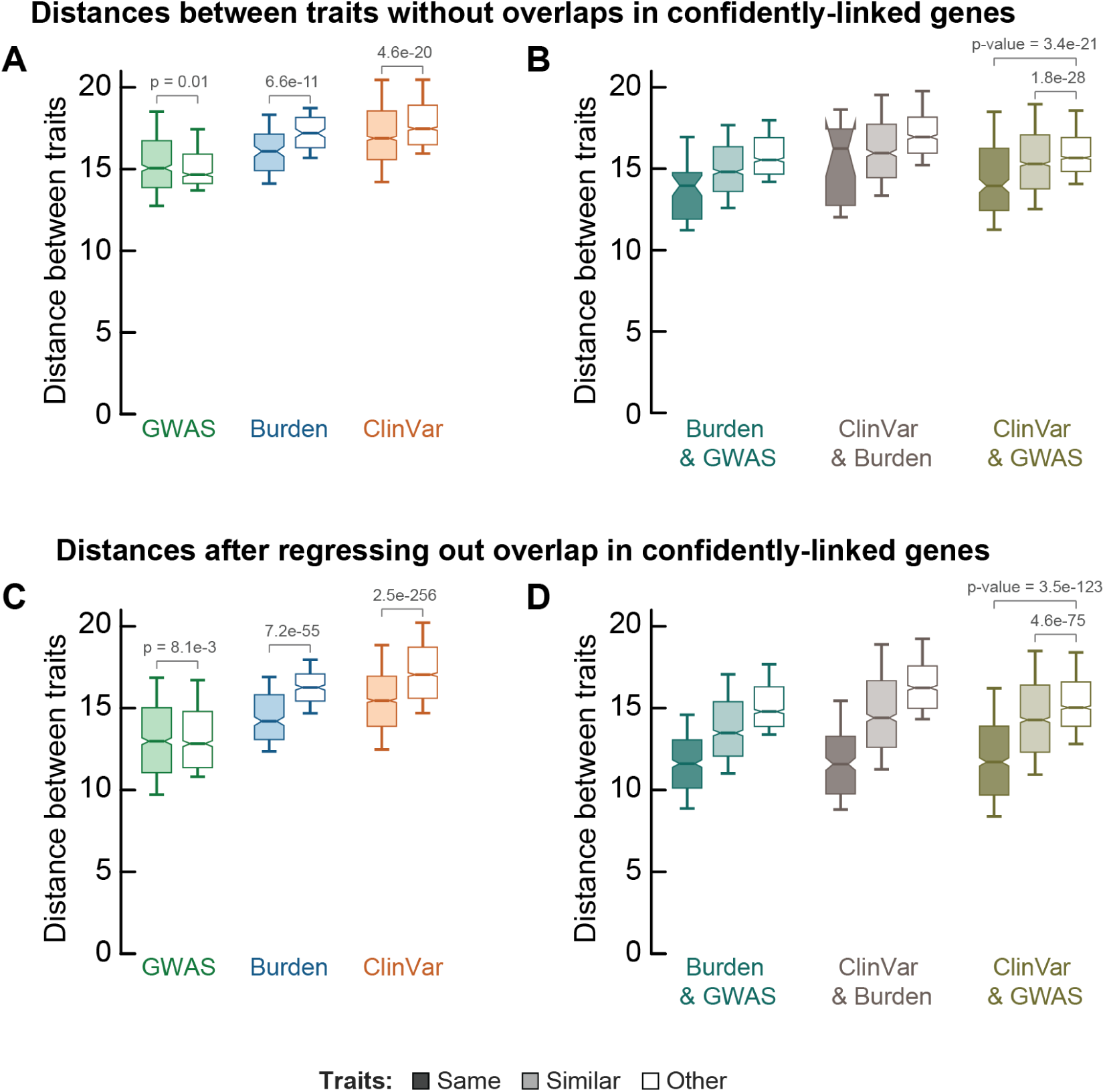
Differences in distances between traits not driven by simple overlaps of linked genes (Related to Fig. 2B**-C**). As expected, we found that the Euclidean distances between latent representations of traits typically correlated with the number of shared confidently-linked genes (ρ = -0.30, p-value = 0.0), although traits typically shared few genes (median jaccard index = 0.01 for the 21% of pairs of traits sharing any genes). We quantified the effect of overlaps in confidently-linked genes on the distances between pairs of traits. **(A-B)** First, we considered pairs of traits that did not share any confidently-linked genes (i.e., analogous to Fig. 2B-C but ignoring pairs of traits with overlaps in trait-linked genes). Shown are relative distances within sources of evidence for GWAS (green), Burden (blue) and ClinVar (orange) **(A)** and across sources of evidence for Burden-GWAS (teal), ClinVar-Burden (brown) and ClinVar-GWAS (dark green) **(B)**. Boxplots show distances between the same traits (dark boxes), similar traits (light boxes) and unrelated traits (white boxes). Traits were filtered for traits having at least 5 confidently linked proteins (c.f. Figs. 2B-C). Similar traits are siblings in the ontology tree, other traits are neither siblings or parent-children (Methods). Boxplots show 10-th and 90-th percentile (whiskers) with the median and s.e. (notches). Metric is the Euclidean distance of latent representations. When only considering the pairs of traits without overlaps in confidently-linked genes, we found that similar traits were still closer compared to unrelated traits for ClinVar (1.05-fold on average; p-value = 4.6e-50; two-sided paired t-test), but not for GWAS (average distances 15.4 and 15.2 respectively; p-value = 0.01). Across sources of evidence, we found that the same trait and similar traits were also still closer compared to unrelated traits (1.14-fold and 1.05-fold respectively; p-values = 3.4e-21 and 1.8e-28 - two-sided paired t-tests). These analyses show that, when only considering pairs of traits without overlaps in linked genes, the same and similar traits are typically still closer in the latent space compared to other traits, even across sources of evidence. **(C-D)** Next, we regressed the overlap in confidently-linked genes out of the distances between pairs of traits. Specifically, we fitted a linear model with intercept to predict the distance between pairs of traits with the Jaccard indices of their confidently-linked genes. The model was fitted on pairs of traits that shared at least one linked gene, and its only parameter was β = − 15. 86 (i.e., distances decrease when the overlap in linked genes increases). We then subtracted the model’s predicted effect of the overlap in linked genes from the distances (i.e., distances between traits without overlaps in confidently-linked genes remained unchanged; distances increased for pairs of traits with overlaps in confidently-linked genes). Shown are these Jaccard-index-adjusted distances within sources of evidence **(C)** and across sources of evidence **(D)** analogous to (A-B) (and for the same pairs of traits as in Figs 2B-C). With the effect of overlaps in confidently-linked genes regressed out of the distances between traits, we found that similar traits were still significantly closer compared to other traits for all sources of evidence (e.g., 1.13-fold; p-value = 2.5e-256 (ClinVar) and 1.03-fold; p-value = 8.1e-3 (GWAS); two-sided paired t-tests) (C). Similarly, across sources of evidence, we found that the same trait was closer compared to other traits (e.g., 1.27-fold; p-value = 3.5e-123 (GWAS-ClinVar)) and that similar traits were closer compared to other traits (e.g., 1.08%; p-value = 4.6e-75 (GWAS-ClinVar)) (D). Together, these results show that differences in distances between related and unrelated traits are not driven by simple overlaps in confidently linked genes, and that similar traits are typically significantly closer compared to unrelated traits even across sources of evidence.

**Figure S4.**
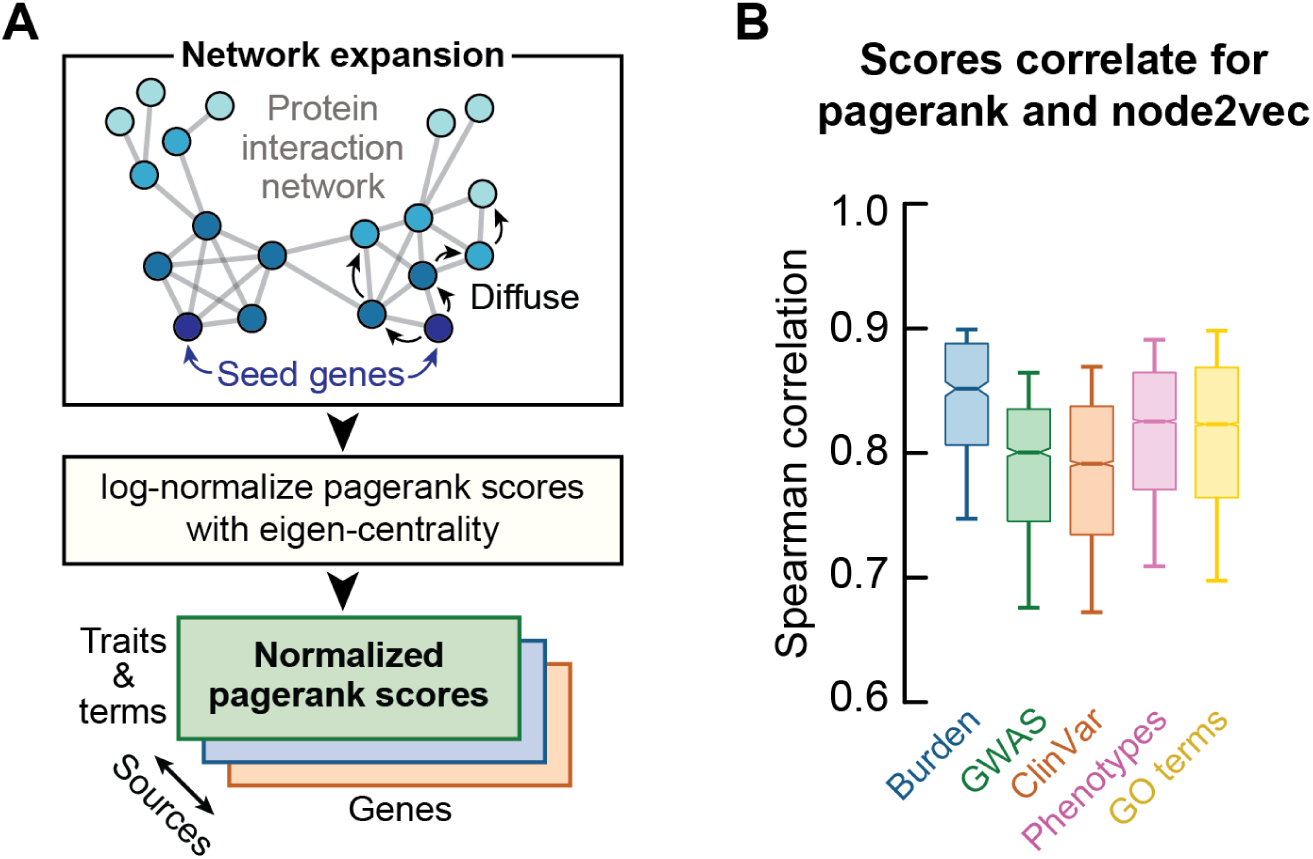
Network scores and latent representations outperform network propagation for separating similar and unrelated traits (Related to Fig. 2B-C). We compared the network scores and their latent representations (Fig. 2A) with the scores obtained from network propagation (pagerank scores). **(A)** For the pagerank scores, we diffused confidently-linked genes for traits and terms over the interaction network using the Personalized PageRank (PPR) algorithm (α = 0.85; implemented from the `networkx` package). These PPR scores for all genes in the network were then log-normalized with the eigencentrality of the nodes in the interaction network to adjust for network biases. With this methodology and following the exact same approach as for the network embedding scores (Methods), we computed pagerank scores for all human traits and terms for each of the sources of evidence, together with the pagerank scores for the sub-samples of traits and terms (Methods). **(B)** We computed the rank correlation of the pagerank and network embedding scores for all terms and traits and all sources of evidence. Boxplots show 10-th and 90-th percentile (whiskers) with the median and s.e. (notches). We found that the pagerank scores strongly correlated with the network embedding scores (median Spearman correlation ρ = 0.82 over all traits and terms), with the scores correlating more strongly for terms (mouse phenotypes (Spearman ρ = 0.83) and GO terms (0.82)) compared to traits from GWAS (0.80) and ClinVar (0.79). We used these PageRank-derived scores to compare with the performance for our proposed network embedding scores using the Node2Vec algorithm.

**Figure S5.**
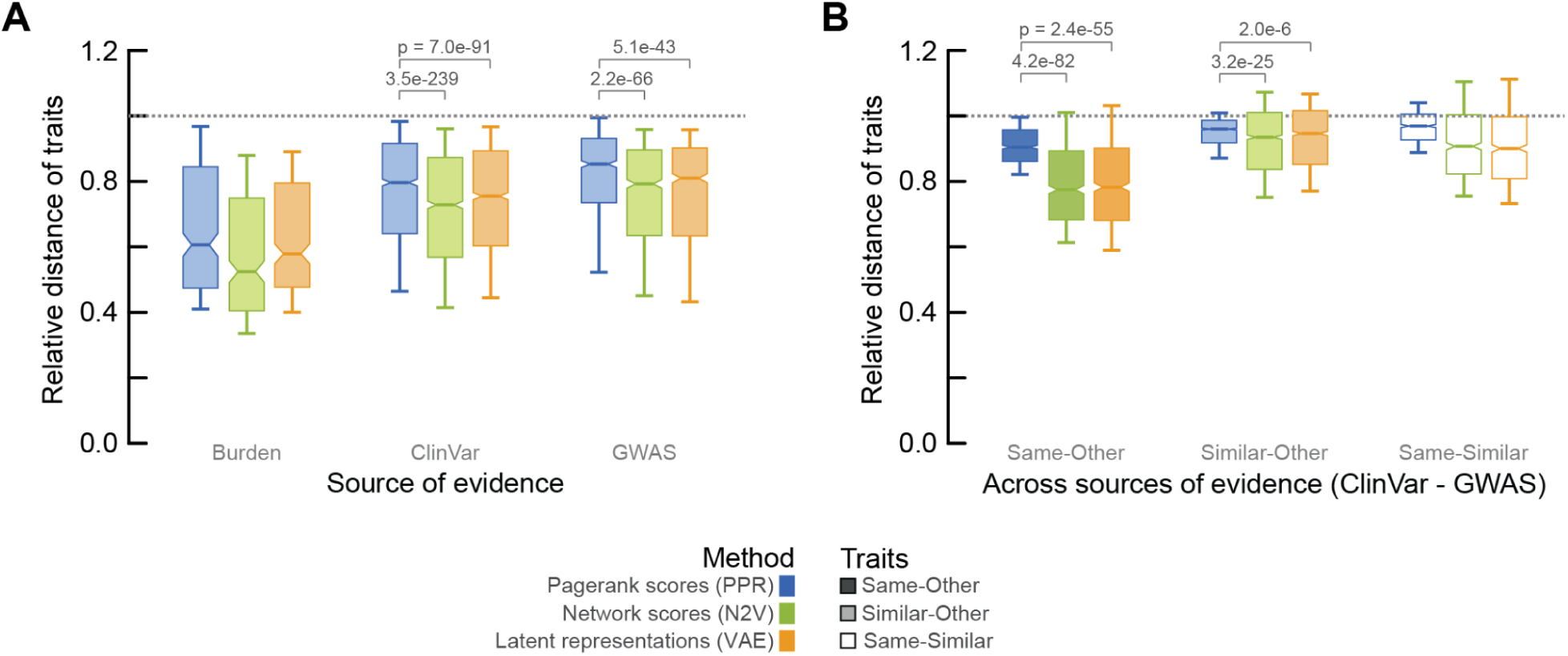
Network embedding scores and latent representations outperform network propagation for separating similar and unrelated traits (Related to Fig. 2B-C). We compared the distances between similar and unrelated traits using the latent representations (c.f. Fig. 2B-C) with distances between traits using the network embedding scores themselves (Fig. 2A) or distances using the PageRank scores (network propagation; Supplementary Fig. 4). To do so, we computed the Euclidean distances between traits using i) the latent representations of the network embedding scores, ii) the network embedding scores, and iii) the normalized pagerank scores. Finally, for each of these methodologies, we compared the distances between traits that were the same, similar (siblings in the ontology tree) or other (not the same, parent-children or siblings - see Fig. 2B-C and Methods). **(A-B)** Shown are the relative distances within **(A)** and across the sources of evidence **(B)**, comparing pagerank scores (blue), network embedding scores (green) and latent representations (orange). For each of these, shown are the relative distances between the same trait relative to distances between similar traits (white), the same traits relative to other traits (dark boxes), and similar traits relative to other traits (light boxes). Distances were first averaged at the trait-level. Boxplots show 10-th and 90-th percentile (whiskers) with the median and s.e. (notches). Compared to the PageRank scores, we found that both the network embedding scores and latent representations of traits improved the separation of similar and unrelated traits for all sources of evidence. For example, the network embedding scores improved the distances of similar relative to other traits by 7.5% for ClinVar (p-value = 3.5e-239; two-sided paired t-test) and 6.7% for GWAS (p-value = 2.2e-66). Similarly, across sources of evidence for ClinVar and GWAS, we found that the network embedding scores improved the distances for the same trait relative to other traits by 12.2% (p-value = 4.2e-82; network embedding scores) and 11.7% (p-value = 2.4e-55; latent representations). Finally, we found that the network embedding scores improved the distances for similar traits relative to other traits by 2.8% (p-value = 3.2e-25), and improved the distances for the same trait relative to similar traits by 4.5% (p-value = 8.0e-19). Together, these results show that the network embedding scores and latent representations outperform more traditional approaches such as network propagation for separating similar and unrelated traits, significantly improving the distances of the same and similar traits relative to other traits within and across sources of evidence.

**Figure S6.**
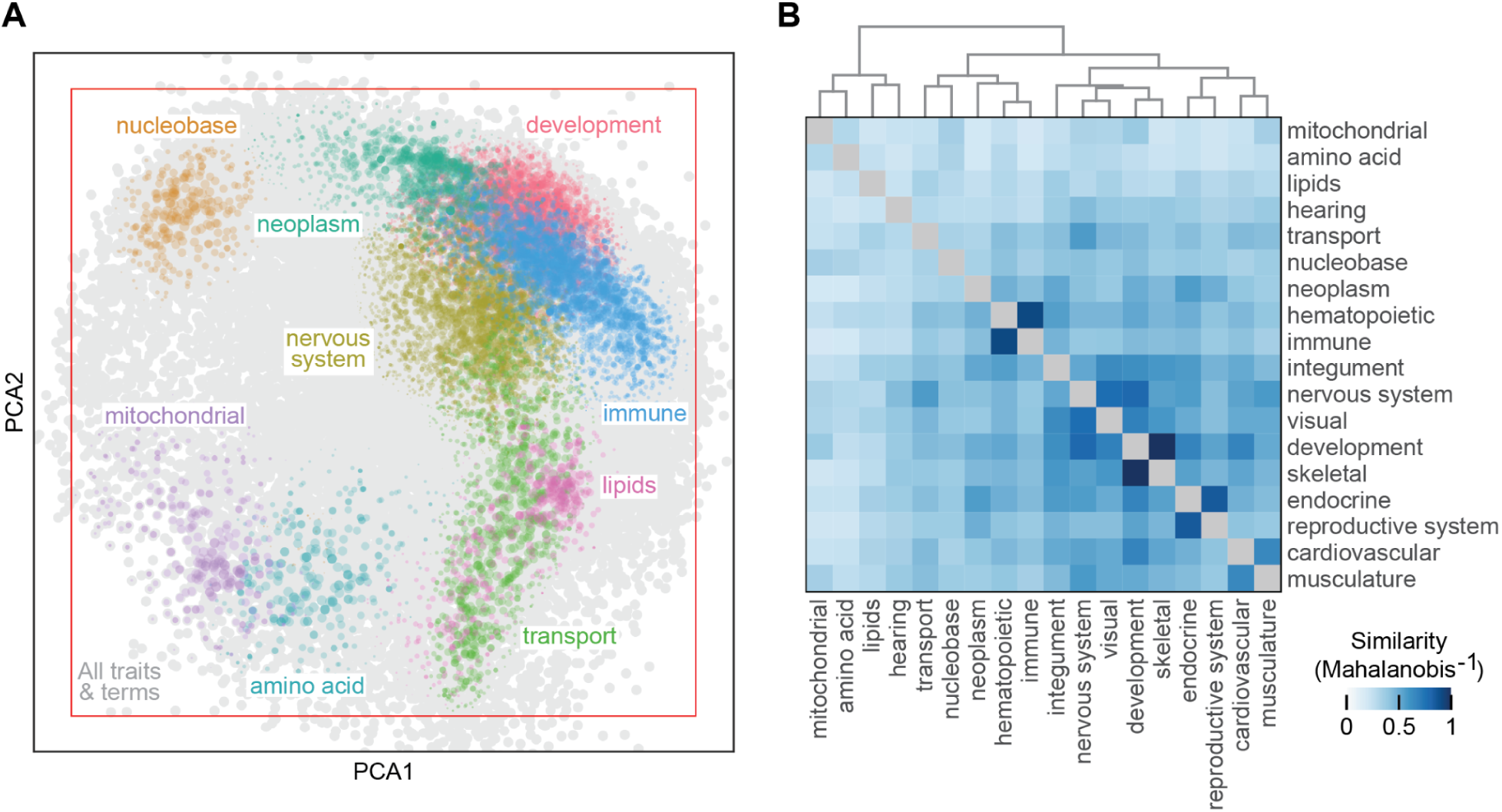
PCA of network scores separates broad categories of traits and terms (Related to Fig. 2D-E). We used the network scores of all 38,826 traits and terms to quantify the explained variance through a Principal Component (PC) analysis. We found that the cumulative explained variance was 0.96 for the first n=128 PCs. **(A)** Scatterplot shows the first two principal components of all traits and terms. Traits and terms were colored according to their most likely broad category from the respective ontology trees (Methods - for selected categories). Probabilities were estimated through a gaussian kernel density for each of the categories. Colored dots were sized proportionally to the probability of the respective category, showing only dots having probabilities exceeding the median probability of all traits and terms (Methods). Background shows all traits and terms (grey). Red square illustrates the approach used for trimming the margins of the UMAP map in Fig. 2D for visualizing traits and terms (Methods). **(B)** Next, we used these principal components to test whether the network scores could separate the broad categories of traits (n=18 categories; Methods; also see Fig. 2c). Heatmap shows the similarity of the n=18 broad categories of terms as quantified through the inverse of the Mahalanobis distances using the first n=128 principal components of the network scores. We found that related categories of terms and traits (e.g., the cardiovascular system and musculature, immune and hematopoietic systems, endocrine and reproductive systems, etc) are more similar (i.e., have a smaller distance) compared to unrelated categories. We further found that 91% of all pairs of categories were significantly different (Benjamini-Hochberg (BH)-adjusted p-values < 0.05) when using only the first principal component of the network scores (explained variance = 0.14; Welch’s t-test), while all pairs of categories were significantly different (BH-adjusted p-values < 0.05) when using the first n>=2 principal components of the network scores (cumulative explained variance >= 0.25; Hotelling’s t2-test - Mahalanobis distance is proportional to the Hotelling’s t2-test statistic). Together, these results show that terms and traits from each of the broad categories group together based on the PCs of their network scores and that the first PCs of the network scores significantly separate the different categories of traits and terms, with related traits and terms having more similar network scores compared to unrelated categories.

**Figure S7.**
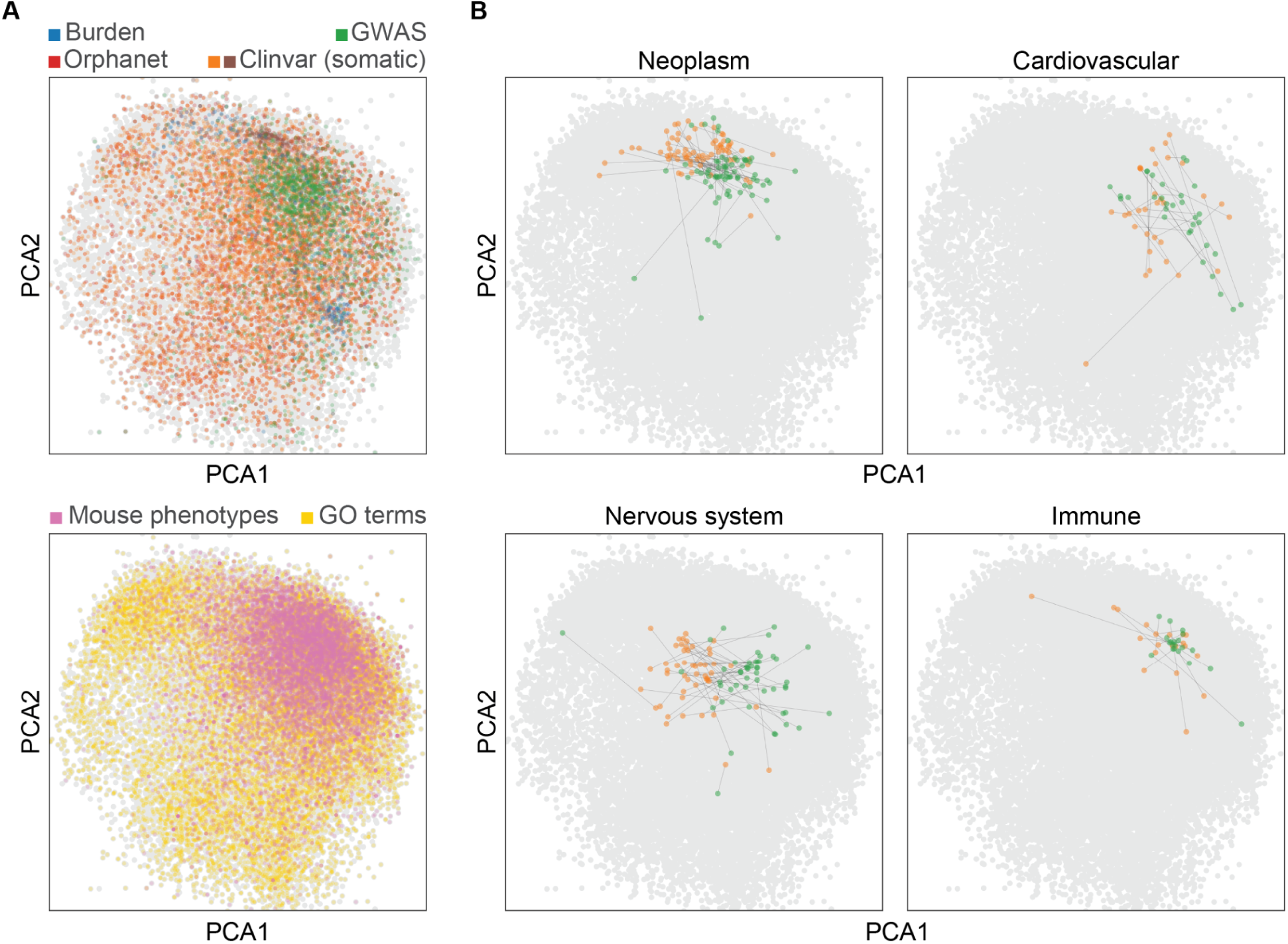
Network scores integrate evidence for terms and human traits (Related to Fig. 2D-E). **(A)** Analogous to Supplementary Fig. 6, we created scatterplots using the first two principal components (PCs) of the network scores for all traits (top) and terms (bottom). Traits and terms were colored by their source of evidence for traits (GWAS, Burden, ClinVar and Orphanet) and terms (bottom - mouse phenotypes and GO terms). All terms and traits are shown as background (grey). **(B)** Next, we selected broad trait categories that had at least 15 traits with network scores from both ClinVar (orange) and GWAS (green). Shown are the scatterplots of the first two PCs of the network scores for neoplasms and diseases of the cardiovascular, nervous and immune systems (visual system not shown). Black lines connect ClinVar (orange) and GWAS (green) for each trait. We found that first two PCs typically separated the same ClinVar and GWAS traits for each of these categories, such as for the nervous system (BH-adjusted p-value = 6.4e-13; Hotelling’s t2-test), but also for all the other sources of evidence (all adj. p-values < 0.05). Together, these examples demonstrate that the network scores integrate the various sources of evidence for traits and terms, with the first two PCs of the network scores separating all pairs of broad trait categories (Supplementary Fig. 6) and the divergent evidence (Fig. 1) underlying source-specific biases within each category of traits.

**Figure S8.**
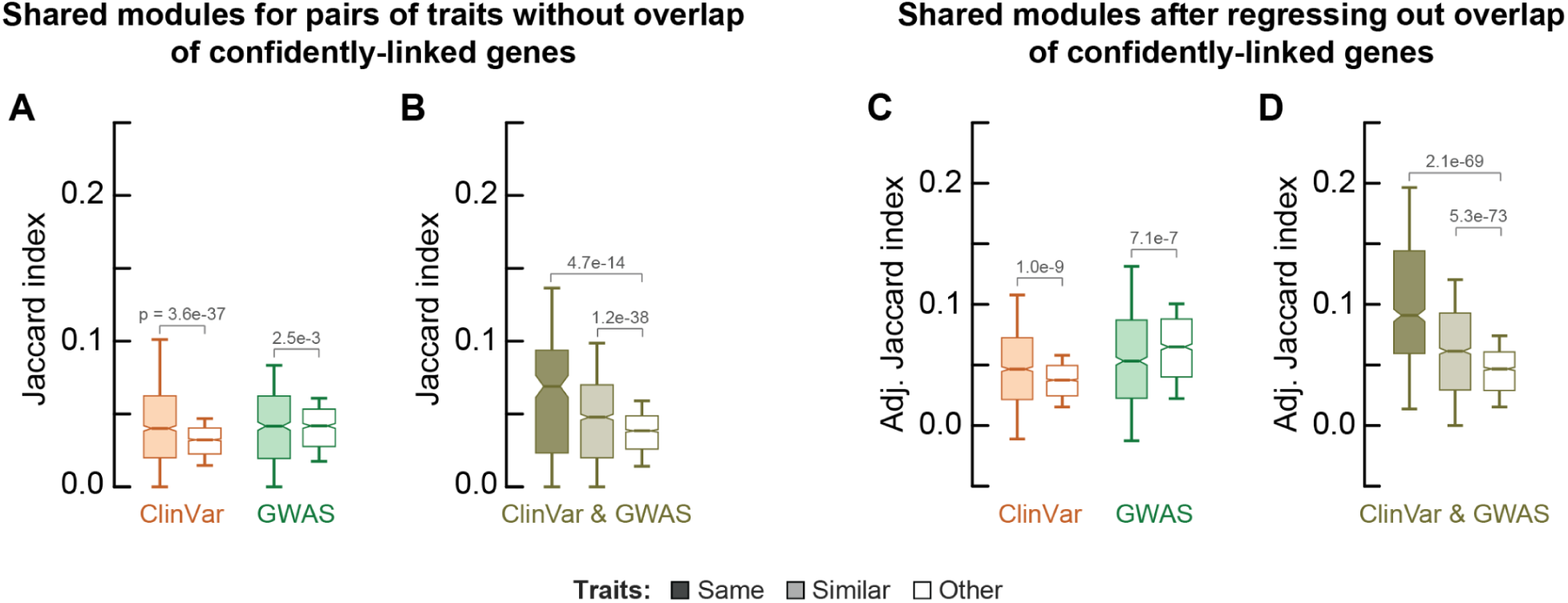
Protein modules shared between traits not driven by simple overlaps of linked genes (Related to Fig. 3B-C). As expected, we found that the protein modules shared between traits correlated with both the overlap in confidently-linked genes (Pearson correlation ρ = 0.67, p-value = 0.0 - Supplementary Fig. 3) and with the latent distances between traits (Pearson correlation ρ = -0.59; p-value = 0.0). As before (Supplementary Fig. 3 - for distances between traits), we quantified the effect of overlaps in confidently-linked genes on the overlaps in linked protein modules between pairs of traits. **(A-B)** First, we considered pairs of traits that did not share any confidently-linked genes. Shown are the shared protein modules as quantified by the Jaccard Index within sources of evidence for GWAS (green) and ClinVar (orange) **(A),** and across sources of evidence for ClinVar-GWAS (dark green) **(B)**. Boxplots show shared modules for the same trait (dark boxes), similar traits (light boxes) and others (white boxes). Traits were filtered for having at least 5 confidently linked proteins (c.f. Figs. 2B-C). Similar traits are siblings in the ontology tree, other traits are neither siblings or parent-children (Methods). Jaccard indices were first averaged at the trait-level. Boxplots show 10-th and 90-th percentile (whiskers) with the median and s.e. (notches). When only considering these pairs of traits without overlaps in confidently-linked genes, we found that similar traits still shared significantly more modules compared to other traits for ClinVar (2.1-fold on average; p-value = 3.6e-37 - two-sided paired t-test) and GWAS (1.1-fold; p-value = 2.5e-3), although similar GWAS traits typically did not share more modules compared to other GWAS traits (median Jaccard indices of 0.417 and 0.418 respectively). Across sources of evidence, we found that the same trait and similar traits shared significantly more modules compared to unrelated traits (1.8-fold and 1.4-fold respectively; p-values = 4.7e-14 and 1.2e-38 - two-sided paired t-tests). These analyses show that, when only considering pairs of traits without overlaps in linked genes, the same and similar traits typically share more modules compared to unrelated traits, especially across sources of evidence. **(C-D)** Next, we regressed the overlap in confidently-linked genes out of the shared protein modules of pairs of traits. Specifically, we fitted a linear model with intercept to predict the Jaccard index of shared protein modules using the Jaccard indices of confidently-linked genes between pairs of traits. Here, the model was fitted on pairs of traits that shared at least one linked gene. The Model’s only parameter was β = 0. 992 (i.e., the number of shared modules increases when the number of shared genes increases). We then subtracted the model’s predicted effect of overlapping genes from the shared protein modules (i.e., the number of shared modules remained unchanged for pairs of traits without overlaps in confidently-linked genes - following Supplementary Fig. 3). Shown are these adjusted Jaccard indices within sources of evidence **(C)** and across sources of evidence **(D)** analogous to (A-B) (and for the same pairs of traits as in Figs. 3B-C). With the effect of overlaps in confidently-linked genes regressed out of the Jaccard index for shared modules between traits, we found that similar traits still shared significantly more modules compared to other traits for ClinVar (1.6-fold; p-value = 1.0e-9 - two-sided paired t-test) but not for GWAS (0.7-fold; p-value = 7.1e-7) (C). Similarly, across sources of evidence (GWAS-ClinVar), we found that the same and similar traits still shared significantly more modules compared to other traits (e.g., 2.1-fold and 1.5-fold respectively; p-values = 2.1e-69 and 5.3e-73). Similar to overlaps in linked genes, we created a linear model to regress the distances in latent representations of traits out of the number of shared modules (Model’s parameter β = -0.01; i.e., shared modules increase as distances decrease as expected). From this model, we found that the latent distances also do not drive the number of modules shared by similar traits exceeding those shared by unrelated traits, both within the sources of evidence (1.7-fold difference for both ClinVar and GWAS; p-values = 3.3e-228 and 6.0e-122 respectively - two-sided paired t-tests) and across sources of evidence (1.19-fold and 1.07-fold (GWAS-ClinVar); p-values 1.9e-49 and 3.8e-57 respectively). Finally, we created a linear model to regress both the overlap in linked genes and the distances from latent representations of traits out of the number of shared modules (Model’s parameters were β_1_ = -0.008 (distances) and β_2_ = 0.90 (shared genes)). From this model, we found that similar traits no longer shared more modules than unrelated traits for any of the sources of evidence (e.g., 0.94-fold for ClinVar). However, across sources of evidence (GWAS-ClinVar) we found that both the same and similar traits still shared more modules than unrelated traits (1.17-fold and 1.07-fold respectively; p-values = 2.4e-36 and 3.1e-45 - two-sided paired t-test). These observations suggest that the overlap in shared genes and latent distances together explain the differences in modules shared between similar and unrelated traits within - but not across - sources of evidence, suggesting that traits from different sources of evidence converge onto functional modules of cell biology that cannot be explained by similarities of their network scores or overlaps in confidently-linked genes. Together, these results show that related traits typically share significantly more modules than unrelated traits, and that these differences across the sources of evidence are not driven by simple overlaps in confidently-linked genes or similarities in their network scores.

**Figure S9.**
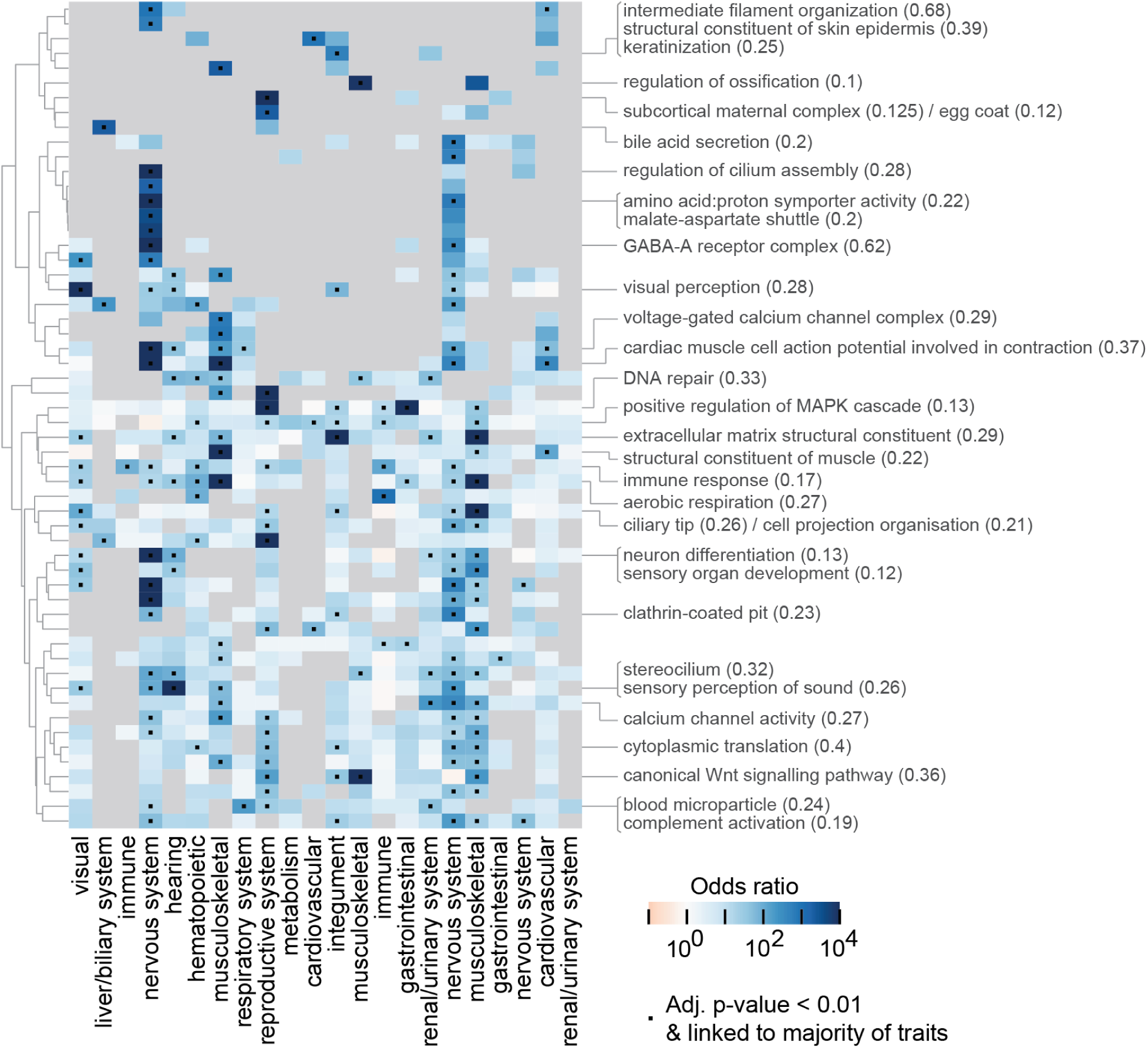
Protein modules linked to clusters of similar ClinVar traits (Related to Fig. 3D). We explored the specificity of functional modules linked to the trait clusters (Fig. 3D). The protein modules were linked to clusters by being associated with at least half the Clinvar traits of the cluster and for being enriched with associations to traits in that cluster compared to other traits that did not share the same annotation (Methods). This enrichment thus links trait clusters to modules in a cluster-specific manner. We selected the protein modules that were linked to clusters from at least three trait categories or that were very strongly linked to at least one trait cluster (odds-ratio > 500; one-sided Fisher exact test - Methods). Shown is the log-odds ratio for the enrichment of associations between the trait clusters (Fig. 3D) and each of these modules. Modules were clustered through complete-linkage clustering with the Manhattan distances of the log-odds ratios; trait clusters are shown in the order following Fig. 2D. Dots show cluster-module links as in Fig. 2E, grey boxes show clusters without any associations with the module. We then labeled the n=56 selected modules through their enrichment with genes from GO terms (one-sided Fisher exact test; Methods). Specifically, modules were labeled with the GO terms having the strongest enrichment for that module (requiring at least odds-ratio >= 10 and BH-adjusted p-value < 0.01; Jaccard indices with the module’s proteins shown in brackets). As expected, we found that modules linked to several categories of traits had more general functions, such as modules for translation, the extracellular matrix, DNA repair, regulation of the MAPK cascade and aerobic respiration (linked to at least 5 clusters from different broad categories of traits). We also found many modules that were very strongly (odds-ratios > 500) and specifically linked to single categories of traits, such as modules for skin epidermis (trait cluster of the integument; BH-adjusted p-value = 1.3e-11; one-sided Fisher exact test (Fig. 3D)), bile acid secretion (liver/biliary traits; 7.8e-8), visual perception (visual; 2.5e-20), stereocilia (hearing; 3.3e-8), GABA-A receptor and neuron differentiation (nervous system; 1e-19 and 2.7e-11), and ossification (musculoskeletal; 1.2e-7) or structural constituent of muscle (musculoskeletal; 5.5e-9). Together, these examples demonstrate that our approach linked modules to trait clusters in a specific and context appropriate manner, identifying specific aspects of cell biology relevant to the diseases.

**Figure S10.**
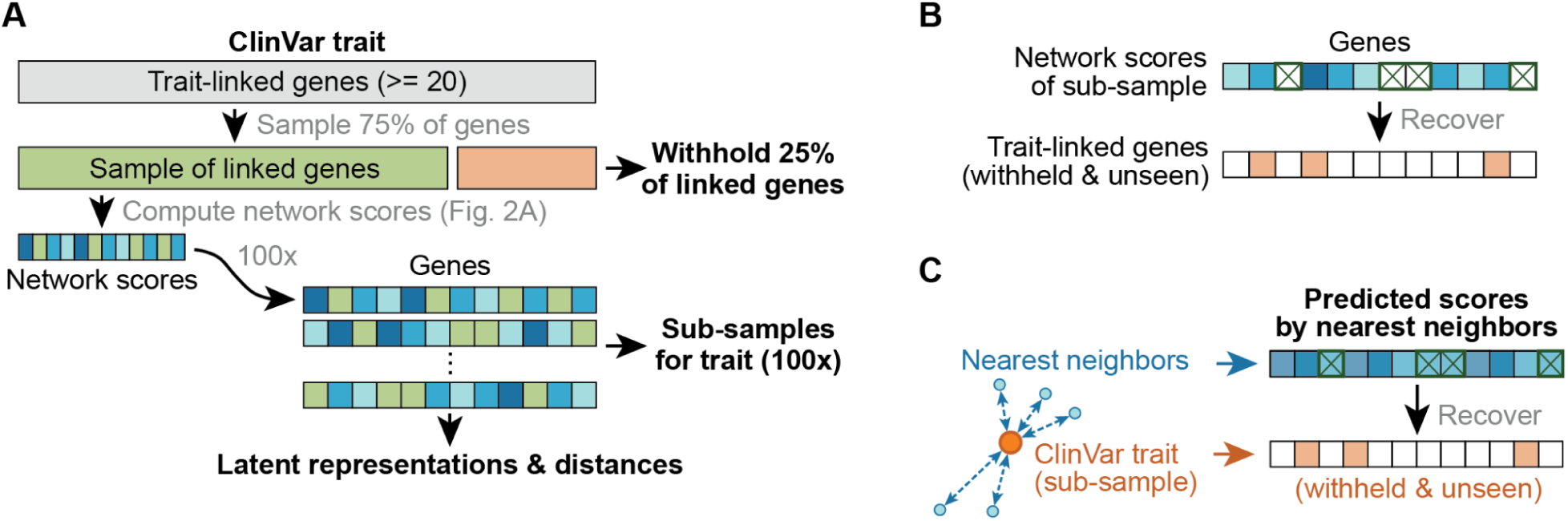
Recovering trait-linked genes using network scores (Related to Fig. 4A-C). **(A)** We used the network scores of sub-samples for each trait (Methods) to recover the trait-linked genes that were withheld for that sub-sample respectively. In short, we selected traits that had at least 20 confidently-linked genes (grey box), and for each trait created n=100 samples with 75% of the trait linked-genes (green box) such that all linked genes were sampled equally frequently and such that each sample withheld 25% of linked genes (orange box - also see Methods). These sampled genes were then used to compute the network scores, latent representations and distances for n=100 sub-samples of each trait (following Fig. 2A). This yielded network scores for sub-samples of 5,303 traits and terms (1,928 GO terms, 2,128 mouse phenotypes, and 704 ClinVar and 543 GWAS traits). Scores were analogously computed for the same samples of traits using the PageRank algorithm (following Supplementary Fig. 4). **(B)** Next, we used the network scores of these sub-samples to recover trait-linked genes (applied in Fig. 4A). Specifically, for each sub-sample, we removed the 75% of trait-linked genes from its network scores (green crosses). The remaining network scores of a trait’s sub-sample were then used as a predictor to recover the 25% of trait-linked genes that were withheld for that sub-sample (i.e., these are known trait-linked genes but are unseen during the process of computing the predicting scores for the sub-sample of the trait). **(C)** Schematic shows approach for recovering trait-linked genes using the nearest neighboring traits or terms. The network scores of the sub-samples are used to select the nearest traits or terms from a given source of evidence (“nearest neighbors”) by some distance metric (Fig. 4C: Euclidean distance in latent space). These nearest neighbors are then used to create predicted network scores (Fig. 4C: averaging their latent representations and then decoding this average with the VAE to obtain reconstructed network scores). Finally, the trait-linked genes from the sub-sample are removed from these predicted scores as before, and the remaining predicted network scores are used to recover the 25% of trait-linked genes that were withheld when computing the network scores for the sub-sample.

**Figure S11.**
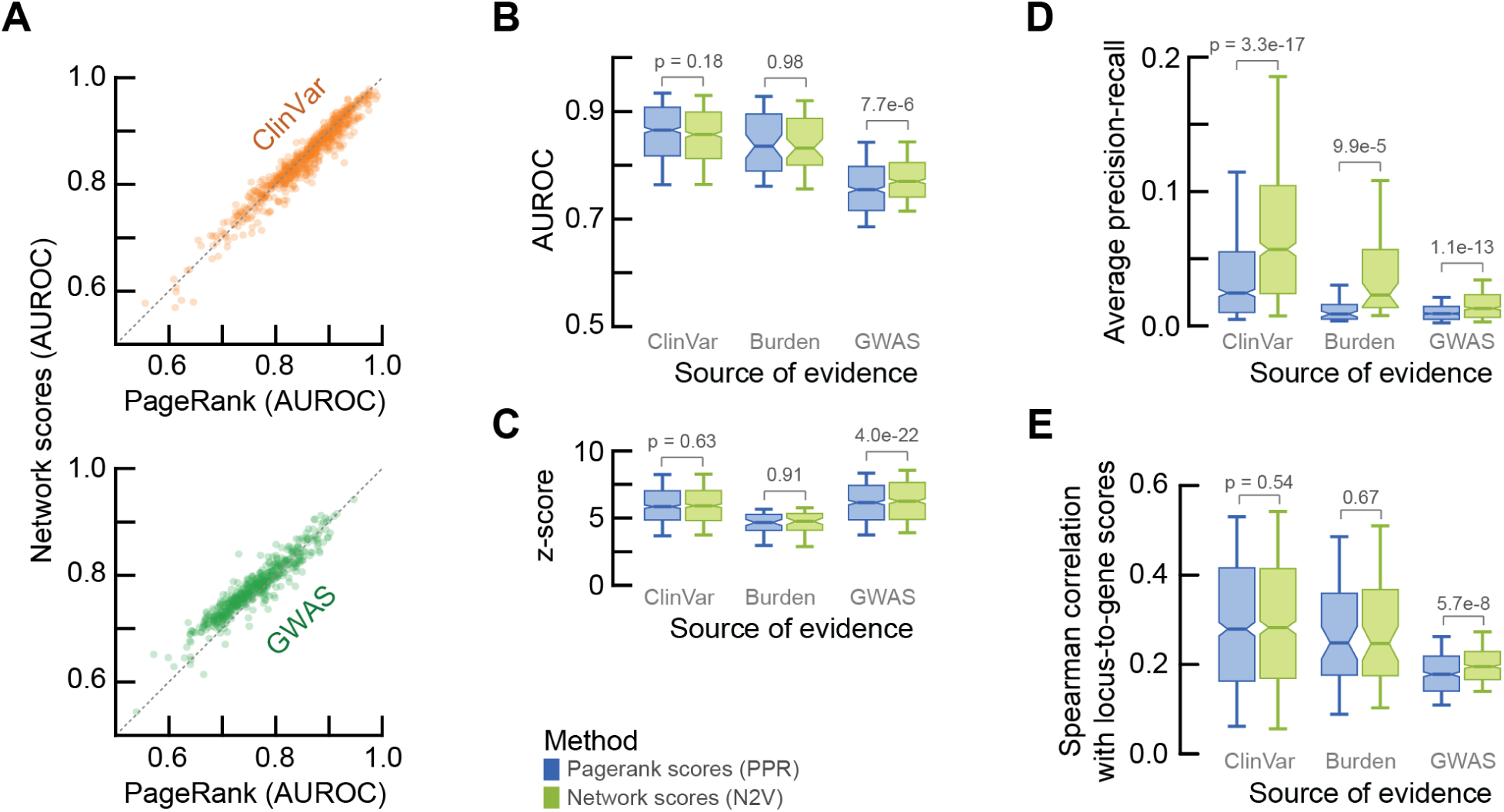
Network embedding scores outperform PageRank for recovering disease-linked proteins (Related to Fig. 4A). We compared the recovery of confidently-linked genes between the network embedding scores and the PageRank scores (Supplementary Fig. 4). Here, we used the network embedding scores for sub-samples of traits that had at least n=20 confidently-linked genes to recover trait-linked genes in a blinded fashion - i.e., using sub-samples of trait-linked genes to predict scores with Node2Vec and PageRank and then recovering the withheld linked-genes from the same trait (Supplementary Fig. 10; Methods). The traits with sub-samples were further filtered for traits whose gene samples from the disease-gene frequency distribution had a median jaccard index at most 0.1 (i.e., sampling the frequency distribution of linked genes from the respective source of evidence yielded sufficiently different samples - see Fig. 4A; Methods). **(A)** Shown is the recovery of the withheld trait-linked genes for the sub-samples of traits with evidence from ClinVar (top) and GWAS (bottom). Each dot represents one trait using scores from PageRank (x-axis) and Node2Vec (y-axis). **(B)** Summarized data from (A), showing the recovery of confidently-linked genes for traits with evidence from ClinVar, Burden and GWAS by scores derived from PageRank (blue) or Node2Vec (green). Aggregated over the traits from all sources of evidence, we found no difference between the recovery of confidently-linked genes by the PageRank scores (average AUROC = 0.814 ± 0.002 (mean with s.e.m.)) or the network embedding scores (0.818 ± 0.002; p-value = 0.20 - two-sided Welch’s t-test). More specifically, for ClinVar traits, we found that the PageRank scores (AUROC = 0.856 ± 0.003) were similar to the network embedding scores (0.851 ± 0.003) for recovering confidently-linked ClinVar genes (p-value = 0.18; two-sided Welch’s t-test). In contrast, for GWAS traits, we found that the network embedding scores (0.774 ± 0.002) outperformed the PageRank scores (0.759 ± 0.003) for recovering confidently-linked GWAS genes (p-value = 7.7e-6). **(C)** For the same traits as in (B), showing the specificity for recovering trait-linked genes compared to other genes sampled from the frequency distribution of linked genes for the respective source of evidence (following Fig. 4a; Methods). Aggregated over the sources of evidence, there were no significant differences in the specificity for recovering trait-linked genes between the scores from PageRank (average z-score = 5.88) and network embeddings (5.95; p-value = 0.35 - two-sided Welch’s t-test). **(D)** Finally, as before (A-C), we computed the average precision-recall (PR) for recovering confidently-linked genes by the scores from PageRank (blue) and network embeddings (green) for ClinVar, Burden and GWAS traits. We found that the PR from the network embedding scores (median PR = 0.057 (ClinVar) and 0.013 (GWAS)) outperformed the PageRank scores (0.024 and 0.009) for all sources of evidence (p-values = 3,3e-17 and 1.1e013 respectively; two-sided Welch’s t-test). Aggregated over the sources of evidence, we found that the network embedding scores improved the average precision-recall by 2.2-fold (p-value = 2.2e-20; two-sided Welch’s t-test). **(E)** Finally, we tested the ability of the network embedding scores to recapitulate the confidence scores from Open Targets (OTAR) for genes to be linked to a trait. We have used confidence scores from OTAR to select the confidently-linked genes for each trait (scores >= 0.5, the same linked-genes were used for both Node2Vec and PageRank; Methods). These trait-linked genes were then sampled to compute the scores for the sub-samples (Methods). Thus, the OTAR confidence scores themselves were not used for computing the network and PageRank scores of the sub-samples. We computed the rank correlations of the network embedding or PageRank scores with the confidence scores from OTAR for genes that had a confidence score and were not used as linked-genes when computing the sub-samples (i.e., network embedding scores and PageRank scores were blinded for the trait-linked genes). Shown are the Spearman correlations after averaging over the sub-samples from each of the ClinVar traits. We found that the network embedding scores correlated more strongly with the OTAR confidence scores compared to the PageRank scores for each source of evidence, although differences were generally small. Indeed, aggregated over the sources of evidence, we found that the correlations were significantly different but generally similar for the network embedding scores (median Spearman correlations of ρ = 0.255) and the PageRank scores (ρ = 0.243; p-value = 0.046 - two-sided Welch’s t-test). This analysis shows that the network embedding scores and PageRank scores correlate with the confidence scores for being linked to traits from Open Targets, with the network embedding scores outperforming the PageRank scores in recapitulating the ranking of genes by Open Targets. Together, these analyses (in A-E) demonstrate that the network embedding scores typically outperform the PageRank scores for recovering confidently-linked genes. However, these differences for recovery of trait-linked genes are generally minor, with some biases between the sources of evidence (e.g, network embedding scores outperforming more strongly for GWAS traits). In (B-E), the p-values on boxplots show significance for two-sided Welch’s t-tests. Boxplots show 10-th and 90-th percentile (whiskers) with the median and s.e. (notches).

**Figure S12.**
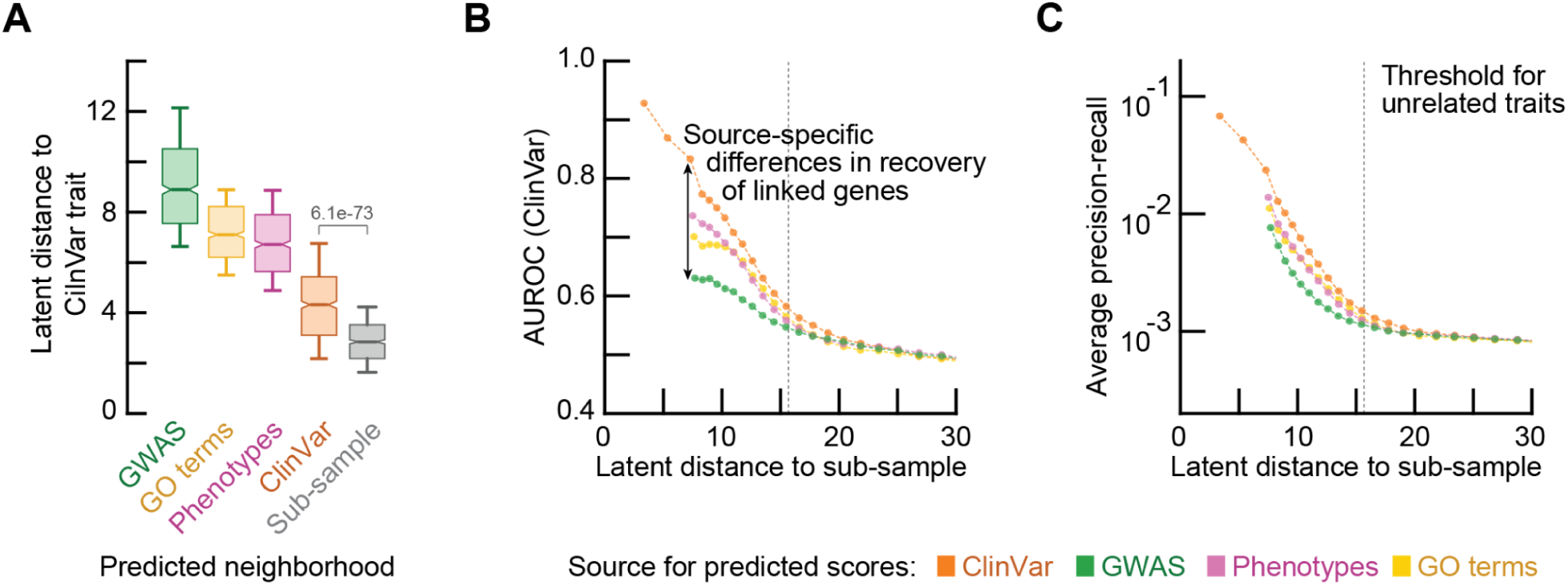
Latent neighbors from ClinVar outperform other sources of evidence for recovering genes linked to rare diseases (Related to Fig. 4C). **(A)** We compared the latent distances of the nearest neighbors from the different sources of evidence. To do so, we selected the n=5 nearest neighbors for each of the sub-samples of the ClinVar traits (i.e., the same nearest neighbors as in Fig. 4C), averaged their latent representations, and then computed the Euclidean latent distance to the respective ClinVar trait. Shown are the latent distances of the ClinVar traits to the averaged nearest neighbors from each source of evidence, compared to the latent distances of the ClinVar traits to its sub-samples. Boxplots show 10-th and 90-th percentile (whiskers) with the median and s.e. (notches). Distances were first averaged over the sub-samples from each of the ClinVar traits. As expected, we found that the ClinVar traits were closest to their sub-samples (average distance = 2.9), followed by the nearest neighbors from ClinVar (distance = 4.4; p-value = 6.1e-73 - two-sided Welch’s t-test). Strikingly, the nearest neighbors from GWAS were on average 2.4-fold further from the ClinVar trait compared to the nearest neighbors from ClinVar (p-value = 8.7e-282). These observations suggest that the recovery of trait-linked genes by the predicted scores from GWAS neighbors may be outperformed by other sources of evidence due to their larger distances to the rare diseases. **(B-C)** To test this, we quantified the recovery of trait-linked genes by neighbors as a function of their latent distance to the trait for each source of evidence. To do so, we computed the distances of all traits and terms with network scores to the first 4 sub-samples of the ClinVar traits (Methods). We then binned these distances on a log-scale for each sub-sample, and for each bin randomly sampled 10 traits and terms per source of evidence when available (ignoring the same ClinVar trait as the sub-sample). Using the network scores for these selected traits and terms, we then computed the recovery of the trait-linked genes that were withheld for each sub-sample (Supplementary Fig. 10; Methods). With this approach, we scored the recovery of genes linked to ClinVar traits as a function of latent distance to the traits and the source of evidence for the predicting neighbor. Finally, we aggregated the scores of the smallest and largest bins if less than 2,000 scores were available per bin, and computed each bin’s midpoint as the median distance of its traits and terms to the sub-samples. Shown are the AUROC **(B)** and average precision-recall **(C)** for recovering trait-linked genes as a function of median distance using traits and terms from ClinVar (orange), GWAS (green), GO terms (yellow) or mouse phenotypes (pink). Each dot represents the median score for one bin. The grey dotted line shows the threshold distance (approx. ∼15.7) that was used beyond which traits were no longer considered to be similar (Fig. 3). As expected, we found that the recovery of trait-linked genes for rare diseases deteriorates as the latent distance of selected neighbors increases. Moreover, traits from any source of evidence perform as a random classifier at distances beyond 25.1 (average AUROC < 0.51 for all sources of evidence). We also found that ClinVar traits significantly outperformed the other sources of evidence for recovering trait-linked genes at the same range of latent distances. Specifically, we found that ClinVar traits (e.g., latent distance = 8.9) outperformed GWAS traits (distance = 8.3; p-value = 1.0 - one-sided Welch’s t-test) for recovering the withheld trait-linked genes (average AUROC = 0.76 and 0.65 respectively). In fact, we found that ClinVar traits could be 1.6-fold more distant compared to GWAS traits (latent distances = 13.5 and 8.3) while achieving a similar performance for recovering the withheld trait-linked genes (AUROC = 0.647 for both sources of evidence; p-value = 0.84 - two-sided Welch’s t-test). These observations demonstrate that the recovery of trait-linked genes decreases for increasingly distant neighbors, and that there are source-specific biases in the network scores that underlie significant differences in recovery of trait-linked genes by traits from different sources of evidence at the same latent distances.

**Figure S13.**
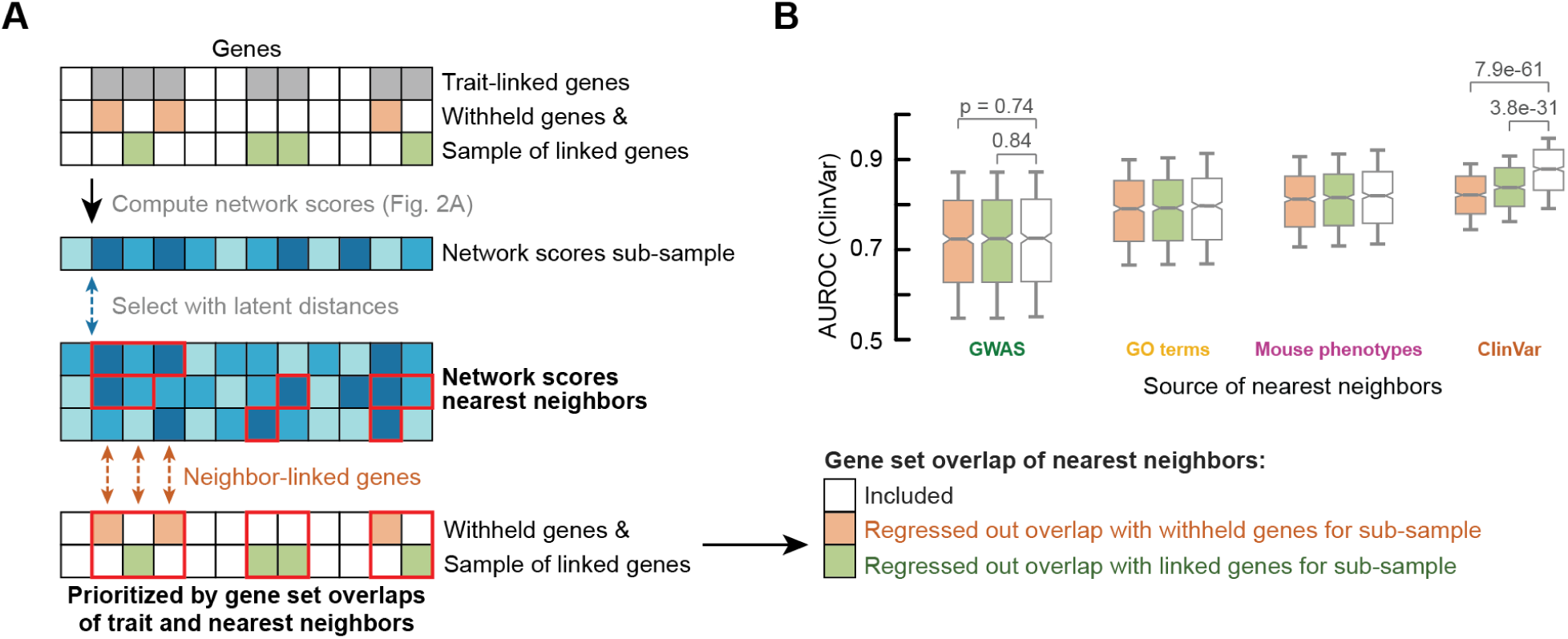
recovery of genes linked to rare diseases not driven by overlaps in trait-linked genes of nearest neighbors (Related to Fig. 4C). **(A)** Sub-samples of genes linked to ClinVar traits were used to compute network scores and used to select the nearest neighbors for the sub-sample through distances between their latent representations. The latent representations of these nearest neighbors were then averaged and decoded to predict network scores and to recover the trait-linked genes withheld for the sub-sample (Fig. 4B-C; Supplementary Fig. 10). As expected, we found that the trait-linked genes of the nearest neighbors in part overlap with the linked genes of the ClinVar trait itself - i.e., the trait-linked genes of the nearest neighbors (red outlines) overlap with the genes used to compute the network scores for the sub-sample (average Jaccard index = 0.321 (ClinVar neighbors) and 0.013 (GWAS neighbors); green boxes) or with the set of withheld genes for the sub-sample (average Jaccard index = 0.125 (ClinVar neighbors) and 0.006 (GWAS neighbors); orange boxes). Since the differences in latent distances between similar and unrelated traits can be in part explained by overlaps in disease-linked genes (Supplementary Fig. 3), we sought to quantify the extent to which the recovery of the withheld genes was driven by these overlaps in disease-linked genes between the rare diseases and their nearest neighbors. **(B)** To do so, we created a linear model to predict the AUCs for recovering the with-held genes of a sub-sample using the predicted scores from the nearest neighbors of the sub-sample. Here, we used as variables the overlaps of the trait-linked genes from each of the n=5 nearest neighbors with either the with-held genes or the trait-linked genes of the sub-sample (i.e., two linear models each having 5 free parameters and an intercept). We then subtracted the part of the AUC that could be explained by the overlaps in linked genes between the neighbors and the sub-sample. Boxplots show the recovery of the withheld genes for the sub-samples (white; reproduced from Fig. 4C), and after regressing out the overlaps between the trait-linked genes of the nearest neighbors from each source of evidence and the trait-linked genes of the sub-sample (orange) or the withheld genes for the sub-sample (green). Boxplots show 10-th and 90-th percentile (whiskers) with the median and s.e. (notches). Scores were first averaged over the sub-samples from each of the ClinVar traits. We found that the overlaps between the trait-linked genes of the nearest ClinVar neighbors and the trait-linked genes of the sub-samples explained a substantial fraction of the recovery of withheld genes (e.g., AUROC = 0.87 compared to 0.84 after regression out overlaps for nearest neighbors from ClinVar; p-value = 3.8e-31 - two-sided Welch’s t-test). Moreover, as expected, the overlap with the (unseen) withheld genes of the sub-samples explained a larger fraction of the recovery of those same withheld genes (AUROC = 0.82; p-value = 7.9e-61). In contrast, the same overlaps in trait-linked genes for the nearest GWAS neighbors did not explain any of the recovery of the withheld genes for the sub-samples (AUROC = 0.72 compared to 0.71 for linked and withheld genes respectively; p-values = 0.84 and 0.74). These analyses show that the predicted network scores from the nearest neighbors of rare diseases recover withheld (unseen) genes beyond simple overlaps between the genes linked to the neighbors and the withheld or linked genes for the rare diseases. The nearest neighbors can therefore be used to prioritize likely casual genes for rare diseases beyond overlaps in linked-genes (i.e., beyond implicating the confident evidence from traits and terms that are close in the latent space).

**Figure S14.**
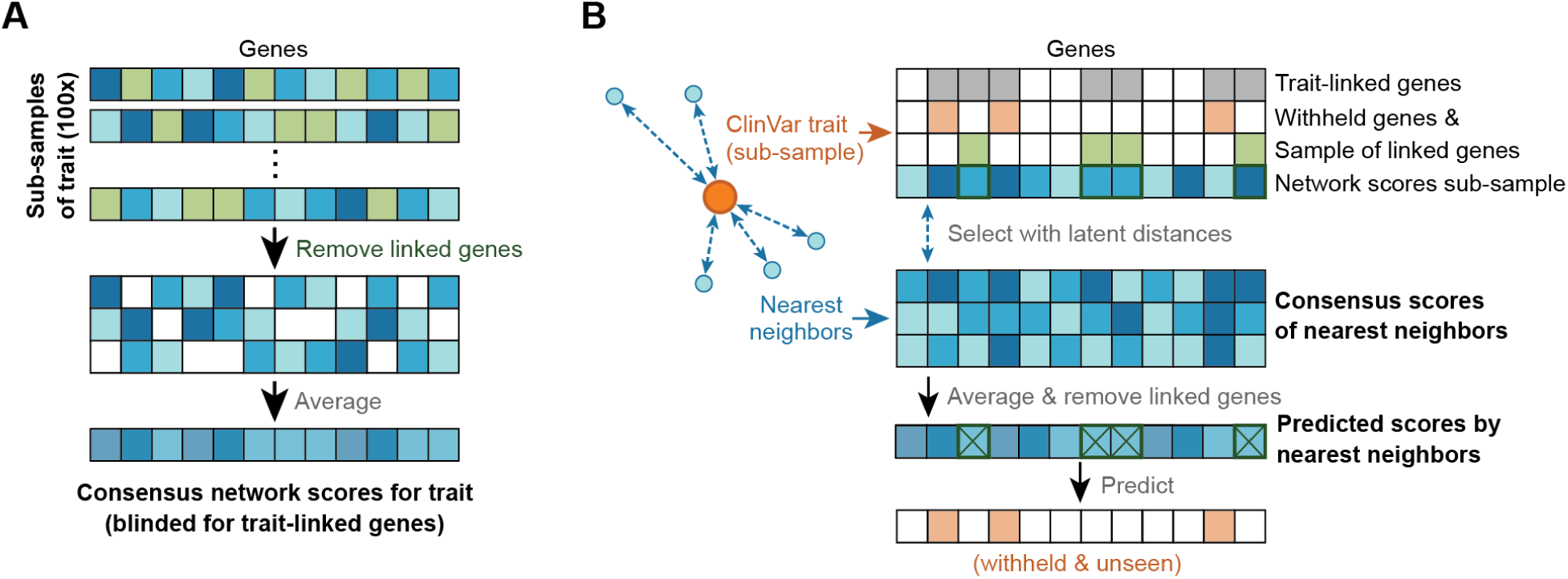
Recovering disease genes with nearest neighbors while blinded for their trait-linked genes (Related to Fig. 4C). We sought to compare the network and PageRank scores for their ability to prioritize genes for rare diseases beyond simple overlaps in trait-linked genes of nearest neighbors. The different algorithms prioritize the known trait-linked genes with different confidence (e.g., average z-score = 2.9 (network scores) and 15.1 (PageRank scores) for the rare diseases), and the genes linked to rare diseases are likely also linked to their nearest neighbors (average Jaccard index = 0.32 (ClinVar) and 0.01 (GWAS); Supplementary Fig. 13). To avoid comparing the performance between the different methods based on overlaps in linked genes, we blinded the predicted scores from the nearest neighbors for their confidently-linked genes. **(A)** To do so, we used the network scores of the sub-samples for each trait (Methods; Supplementary Fig. 10) to create “consensus scores” for the trait that are blinded for the trait-linked genes that were used for each sub-sample to compute its network scores. Specifically, for each sub-sample of a trait, we removed the 75% of trait-linked genes from its network scores and then averaged the remaining network scores over the 100 sub-samples into a consensus score for the trait (i.e., each gene’s consensus score is the average network score of the sub-samples that did not use that gene as a confidently-linked gene). Overall, this approach for computing consensus network scores yields a score for all genes in the interaction network while blinded for the trait-linked genes that were used to compute the network scores. This approach was useful, as it enables the creation of latent representations. Specifically, the VAE does not permit input with missing values, preventing the creation of latent representations for the individual sub-samples when blinding their network scores for their trait-linked genes. The consensus network scores were encoded with the trained VAE using the output of the ‘mean’ layer from the encoder (Methods). Consensus scores were analogously computed for the same traits using the same sub-samples of genes with the PageRank algorithm (Supplementary Fig. 4). In total, we created consensus network scores for n=5,303 traits and terms (1,928 GO terms, 2,128 mouse phenotypes, and 704 ClinVar and 543 GWAS traits). **(B)** Next, we used latent distances for these consensus network scores to select the nearest neighbors for the sub-samples of rare diseases, then decoded the consensus scores of the nearest neighbors to predict scores and recover the trait-linked genes that were withheld for the sub-sample (following Supplementary Fig. 10). Overall, this approach creates consensus scores for a trait that are blinded for its trait-linked genes and can be used to create latent representations and predict network scores from nearest neighbors in a blinded fashion.

**Figure S15.**
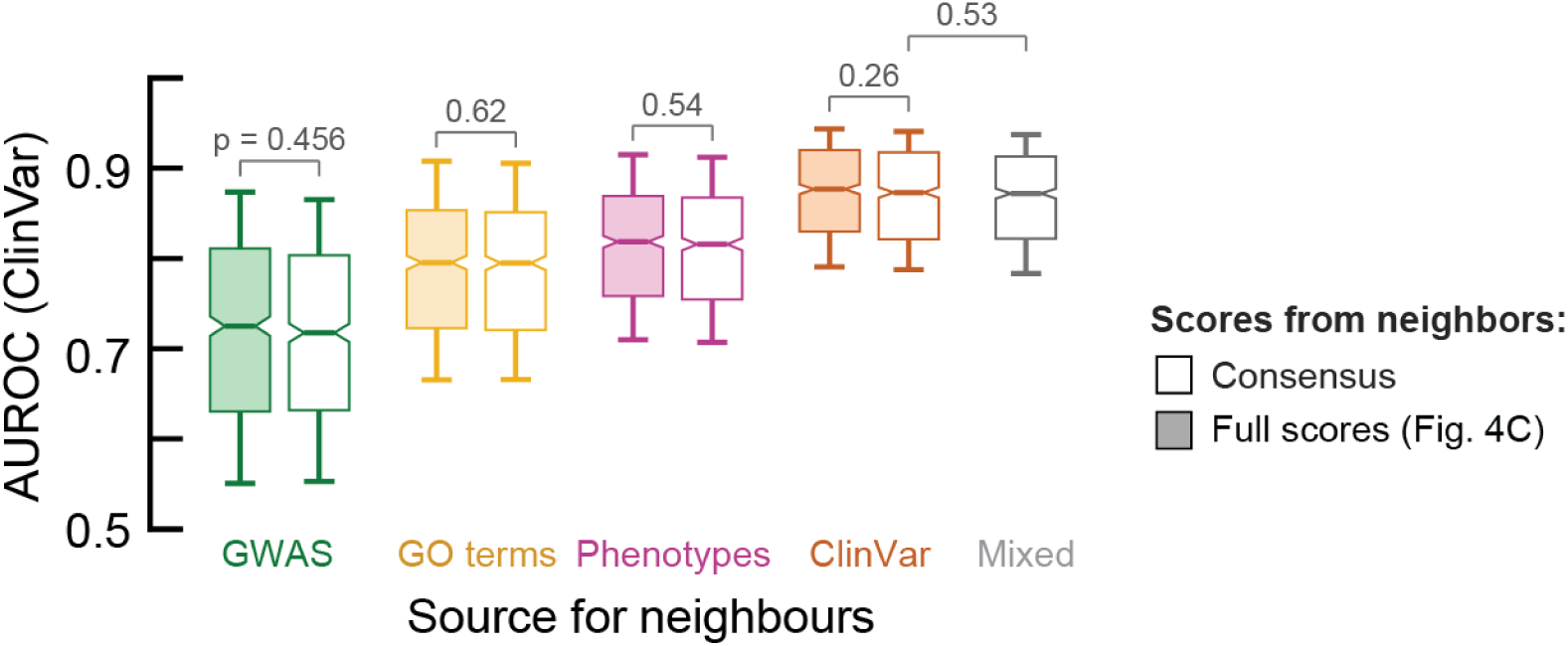
Nearest neighbors from mixed sources of evidence do not improve recovery of trait-linked genes for rare diseases (Related to Fig. 4C). We reproduced the recovery of trait-linked genes using the nearest neighbors for the rare diseases (Fig. 4C) using the consensus network scores (Supplementary Fig. 14). Specifically, we used the latent representations of the consensus scores for n=5,303 traits and terms (Supplementary Fig. 14) to select the nearest neighbors for each of the first n=24 sub-samples of the ClinVar traits through their latent distances. We then predicted network scores from the n=5 nearest latent neighbors for each sub-sample, by averaging and then decoding the latent representations of their consensus scores. With this approach, the predicted scores were blinded for both the genes confidently-linked to the nearest neighbors and for the trait-linked genes withheld for the respective ClinVar sub-sample (Supplementary Fig. 14). Shown is the recovery of the withheld linked genes for the sub-samples using the predicted scores from the nearest neighbors, i) using the consensus scores for each of the sources of evidence (open boxes - analogous to Fig. 4C), ii) using the consensus scores for neighbors from any source of evidence (“mixed sources” - open grey box), and iii) using the network scores as before (colored boxes - reproduced from Fig. 4C; filtered for the same sub-samples). Boxplots show 10-th and 90-th percentile (whiskers) with the median and s.e. (notches). Scores were first averaged over the sub-samples from each of the ClinVar traits. We found no substantial differences between using the network scores or the consensus scores for the nearest neighbors (AUROC = 0.797 and 0.794 respectively; p-value = 0.24 - two-sided Welch’s t-test), even for specific sources of evidence such as ClinVar (0.872 and 0.867 respectively; p-value = 0.26) and GWAS (0.715 and 0.710; p-value = 0.46). Indeed, the network scores and consensus scores strongly correlated (Pearson ρ = 0.99), and minor differences between the scores may be explained between the different sets of traits and terms available to be selected as nearest neighbors. Together, these observations demonstrate that the recovery of trait-linked genes through the nearest latent neighbors of rare-diseases is not driven by the prioritization of genes confidently linked to these neighbors. Finally, we found no substantial differences between selecting the nearest neighbors from any source of evidence (average AUROC = 0.868) or using the nearest neighbors from specific sources such as ClinVar (AUROC = 0.870; p-value = 0.53; two-sided Welch’s t-test). Indeed, we found that the recovery of trait-linked genes strongly correlated between these different predicted network scores (e.g., Spearman ρ = 0.98 between the AUCs from ClinVar or the mixed sources). These results demonstrate that combining the nearest neighbors from any source of evidence does not improve the recovery of trait-linked genes for the rare diseases. This is supported by the different sources of evidence prioritizing different aspects of cell biology (Fig. 3) and our previous observation that there are substantial differences in the recovery of trait-linked genes by traits at the same latent distance but from different sources of evidence, with ClinVar significantly outperforming the other sources of evidence (Supplementary Fig. 12).

**Figure S16.**
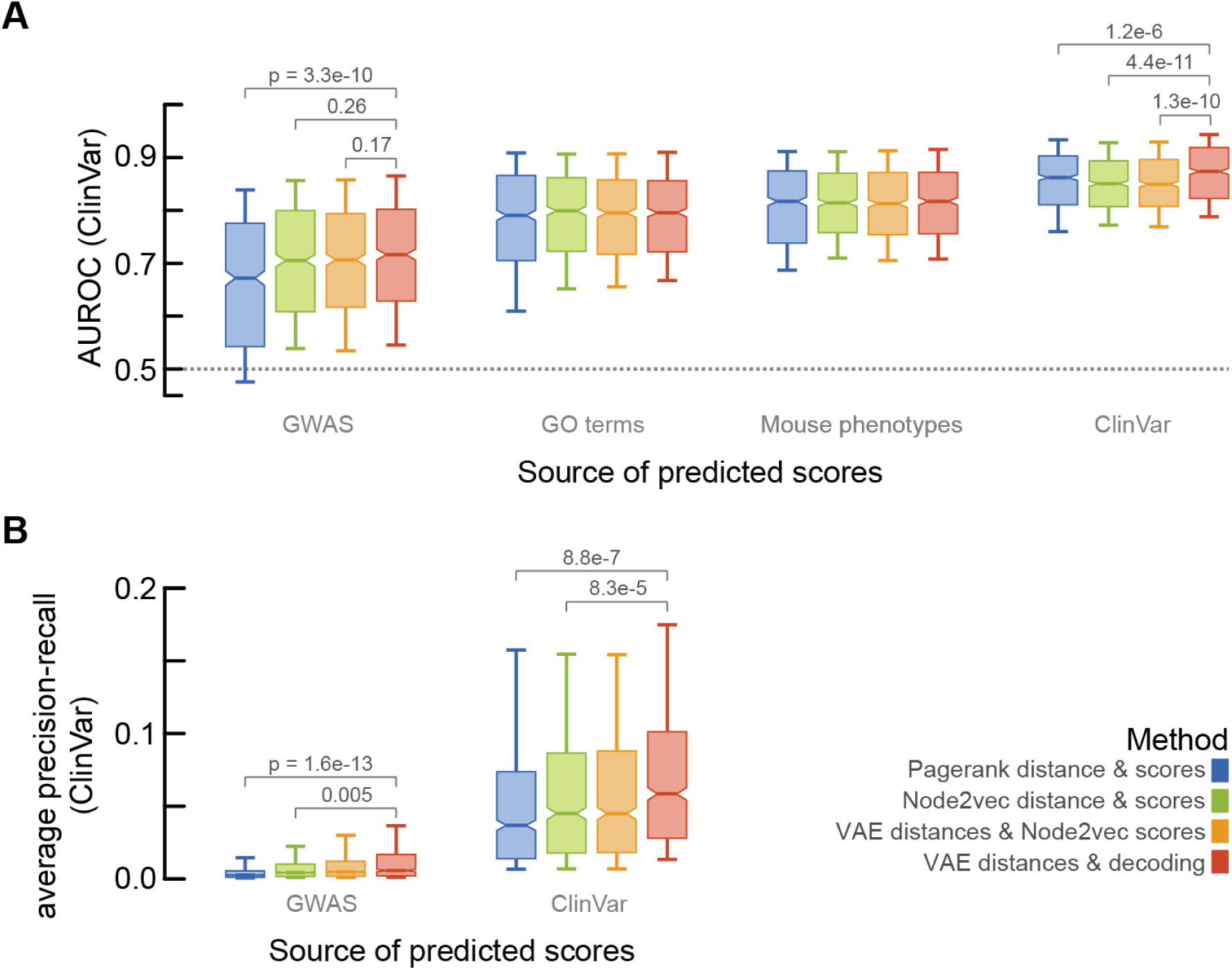
Network embedding scores and latent representations improve recovery of disease-linked genes between traits and across sources of evidence (Related to Fig. 4C). We compared the network embedding scores and PageRank scores (Supplementary Fig. 4) for their performance in recovering trait-linked genes using the predicted scores obtained from the nearest neighbors. To do so, we used the first n=24 sub-samples for each of the n=704 ClinVar traits with at least 20 confidently-linked genes (Methods). For each of these sub-samples, we then selected the nearest neighbors from a given source of evidence, as quantified through some distance metric. For these nearest neighbors, we computed the distances and predicted scores using their consensus scores (Supplementary Fig. 14). Specifically, we used the consensus scores for n=5,303 traits and terms to select the top n=5 nearest neighbors from each source of evidence for the sub-samples of the ClinVar traits, and then recovered the trait-linked genes that were withheld for each sub-sample respectively (Methods). No distance threshold was applied for nearest neighbors (in contrast to Fig. 4) due to differences in distance metrics between the various approaches. **(A-B)** Shown is the recovery of the withheld trait-linked genes by the nearest neighbors from the various sources of evidence as quantified through the AUROC **(A)** and average PR **(B)**. Shown is the recovery for i) averaging and then decoding the latent representations of the nearest latent neighbors (red - using Euclidean distances between latent representations), ii) averaging the network embedding scores of the nearest latent neighbors (orange), or averaging the iii) network embedding scores (green) or iv) PageRank scores (blue) of the nearest neighbors (both using Euclidean distances between the scores themselves). Boxplots show 10-th and 90-th percentile (whiskers) with the median and s.e. (notches). Scores were first averaged over the sub-samples from each of the ClinVar traits. Aggregated over the sources of evidence, we found that averaging and decoding the nearest latent neighbors (average AUROC = 0.794; average PR = 0.048 - red) outperformed averaging the PageRank scores of nearest neighbors (0.774 and 0.036; p-values = 1.6e-10 and 1.4e-12 respectively; two-sided Welch’s t-tests - blue), averaging the network embedding scores of nearest neighbors (0.785 and 0.040; p-values = 7.1e-3 and 1.9e-6 - green), and averaging the network embedding scores of the nearest neighbors in latent space (0.786 and 0.044; p-values = 1.7e-3 and 7.1e-3 - orange). Moreover, the PageRank scores were also outperformed by the network embedding scores when simply averaging the scores from the nearest neighbors (p-value = 8.2e-5; two-sided Welch’s t-test). Looking more specifically at the common and rare variant studies, we found that averaging and then decoding the nearest latent neighbors (blue) outperformed averaging the PageRank scores of nearest neighbors for both ClinVar (AUROC = 0.873 and 0.862 respectively - p-value = 1.2e-6; two-sided Welch’s t-test) and GWAS (0.716 and 0.672; p-value = 3.3e-10). Together, these observations demonstrate that the network embedding scores typically outperform the PageRank scores for recovering trait-linked genes using the nearest neighbors of rare diseases - especially when using latent distances and decoding the latent neighbors to obtain reconstructed network scores - although the absolute differences are generally minor. The p-values on the box-plots in (A-B) show significance for two-sided paired t-tests.

